# Dynamic, adaptive sampling during nanopore sequencing using Bayesian experimental design

**DOI:** 10.1101/2020.02.07.938670

**Authors:** Lukas Weilguny, Nicola De Maio, Rory Munro, Charlotte Manser, Ewan Birney, Matt Loose, Nick Goldman

## Abstract

One particularly promising feature of nanopore sequencing is the ability to reject reads, enabling real-time selection of molecules without complex sample preparation. This is based on the idea of deciding whether a molecule warrants full sequencing depending on reading a small initial part. Previously, such decisions have been based on *a priori* determination of which regions of the genome were considered of interest. Instead, here we consider more general and complex strategies that incorporate already-observed data in order to optimize the rejection strategy and maximise information gain from the sequencing process. For example, in the presence of coverage bias redistributing data from areas of high to areas of low coverage would be desirable.

We present BOSS-RUNS, a mathematical and algorithmic framework to calculate the expected benefit of new reads and generate dynamically updated decision strategies for nanopore sequencing. During sequencing, in real time, we quantify the current uncertainty at each site of one or multiple reference genomes, and for each novel DNA fragment being sequenced we decide whether the potential decrease in uncertainty at the sites it will most likely cover warrants reading it in its entirety. This dynamic, adaptive sampling allows real-time focus of sequencing efforts onto areas of highest benefit.

We demonstrate the effectiveness of BOSS-RUNS by mitigating coverage bias across and within the species of a microbial community. Additionally, we show that our approach leads to improved variant calling due to its ability to sample more data at the most relevant genomic positions.

## 1 Introduction

Third-generation sequencing provides the unprecedented ability to generate reads consisting of multiple kilo- or even megabases (Payne et al. 2019) compared to previous approaches which are limited to sequencing fragments of several hundred bases (after which the error rate increases significantly: Tan et al. 2019). This technological advance has critical implications for many applications in genomics. For example, ultra-long reads are highly useful in increasing assembly contiguity and even allow the construction of telomere-to-telomere assemblies of entire chromosomes (Jain et al. 2018; Miga et al. 2020). Moreover, they can be used to interrogate variation in regions of a genome that are hard to decipher, such as repeats, centromeres or segmental duplications (Shafin et al. 2021), or to generate fully-phased chromosome-level epigenetic maps (Lee et al. 2020).

One way of generating long sequencing reads is through the use of nanopores. This concept was first explored in the 1980s and commercialized in 2014 by Oxford Nanopore Technologies (ONT; Deamer et al. 2016). It relies on the idea of using a protein nanopore as a biosensor that measures the fluctuations of an ionic current across the pore caused by the presence of nucleotides of a translocating DNA or RNA molecule.

Besides providing long reads, ONT’s instruments offer several additional advantages. First, single-molecule real-time sequencing of molecules is possible without the need for prior amplification and can also be used to directly read RNA without reverse transcription (Garalde et al. 2018). The generation of sequencing reads in real time, which in combination with fast library preparation immensely reduces the time needed to go from biological sample to data analysis, enables (e.g.) intraoperative decision making (Djirackor et al. 2021), improved global food security by rapid identification of plant viruses (Boykin et al. 2018) and portable genomic surveillance (Quick et al. 2016).

The most notable drawback of sequencing using nanopores is the elevated rate of sequencing errors compared to previous technologies. However, ONT’s nanopores, sequencing chemistry and basecallers, which are used to translate electrical signal into nucleobases, have been steadily improved and initial error rates of almost 40% (Goodwin et al. 2015) have decreased to as low as ~1% (Sereika et al. 2021) approaching the accuracy of >99% offered by short-read platforms.

A particularly exciting and unique feature of sequencing using nanopores is the ability to reverse the voltage across the pores in order to reject fragments before reading them in their entirety, termed adaptive sampling or “Read Until” in its original implementation (Loose et al. 2016; Oxford Nanopore Technologies 2020). This enables real-time selection of molecules during sequencing based on a small initial part of a read without the need for complex sample preparation. Initially, identifying the genomic origin of these small fragments was achieved by matching the electrical signal (‘squiggles’: Loose et al. 2016) directly to reference genomes translated into simulated current traces. Recent improvements, however, harness the computing power of GPUs for real-time basecalling, making it possible to utilize optimized bioinformatics tools for further processing, e.g. for alignment of reads to reference genomes in base-space (Payne et al. 2021). This has led to much interest in experiments that can be aided by real-time selection of molecules for sequencing (e.g. Miller et al. 2020; Patel et al. 2021; Marquet et al. 2022; Stevanovski et al. 2022).

In current implementations, underlying decisions about which reads to sequence completely are based on *a priori* choice, e.g. of regions of interest (ROIs) in a sequenced genome (Loose et al. 2016; Payne et al. 2021). This restricts their application to a narrow range of problems, where considerable background information is available in advance of sequencing a (potentially poorly characterised) sample. We propose that, in addition, such decisions could also incorporate the information obtained from already-sequenced fragments generated in the current sequencing run. This would allow for optimised, dynamic decisions that maximise the information gain during sequencing, leading to various potential advantages such as reduced time-to-answer, reduced cost, and increased confidence in variation calling.

More specifically, during a sequencing experiment we might observe a distribution of coverage depth that does not correspond well to the aim of the experiment (observed vs. ideal coverage in Fig. 1A, B). For example, for resequencing or variant calling most positions in a genome will match a known reference and their genotype can therefore be confirmed by observing only few reads. In contrast, some sites or regions will be different and their detection and identification might be the goal of such experiments. At these sites we ideally want to sample more sequencing data. Commonly at present, the overall coverage of the target genome would have to be increased in order to ensure sufficient sampling in biased regions. This leads to wasteful data acquisition in regions that are not of continued interest. In this work we address this issue by generating dynamic decision strategies that redistribute sequencing coverage to positions of greatest value at any point during an experiment.

**Figure 1:**
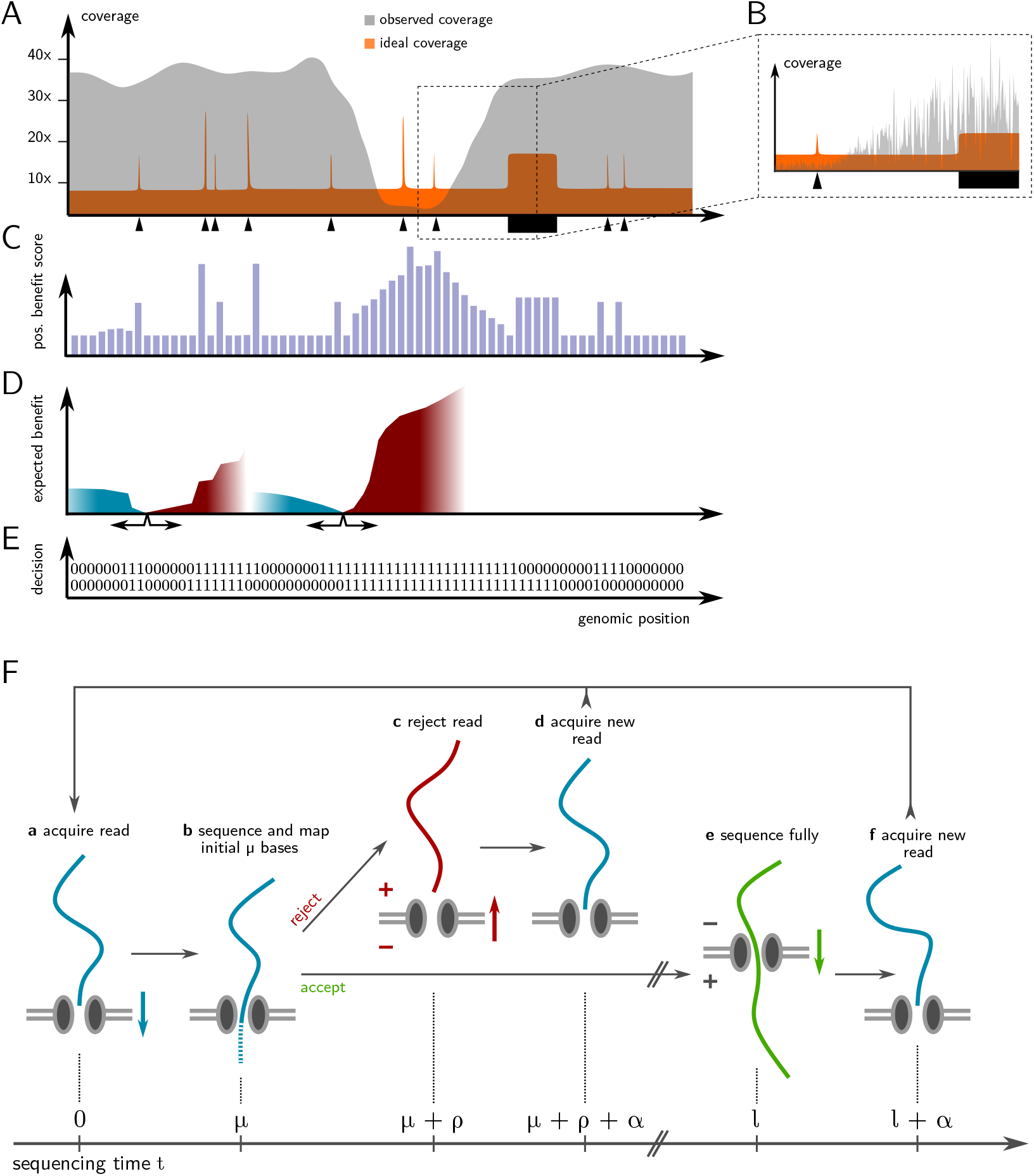
Methodological overview. A) When re-sequencing a genome, different sites might require different levels of coverage. For example, sites without variation are resolved by few reads (low ideal coverage). Additional accumulation of coverage at those positions is wasteful, whereas other sites with higher uncertainty (e.g. due to the presence of variation or lower-quality reference genome, marked by black triangles and rectangles) would benefit from more observations. B) Another reason for reduced efficiency of sequencing could be local fluctuations in the distribution of fragments’ origins, resulting in uneven coverage. C) We quantify the uncertainty about the genotype at each site by calculating posterior probabilities of all possible genotypes, based on prior probabilities and the sequence data observed so far. The expected shift in genotype uncertainty caused by observing a new read at that position is expressed as a “positional benefit score”. D) Next, we calculate the total expected benefit from a hypothetical read starting at each location, i.e. the sum of the positional scores that such a novel read might accumulate (depicted by the height of the expected benefit plots), weighted by the probability of reaching those positions (depicted by the color-gradients). As an example, the benefit of both forward (red) and reverse-oriented reads (blue) starting at two positions are shown. E) Finally, we derive a decision strategy for each position, shown as zeros and ones, that instructs the sequencer to either continue sequencing (1) or reject from the pore (0) a read that starts at that position. Stages C–E can be updated and iterated throughout the sequencing experiment. F) Overview of our model of the sequencing process. Since we parameterize time by the speed of bases translocating through pores we use number of bases and sequencing time interchangeably. At first (*t* = 0), a novel read is acquired by a nanopore and sequencing commences. After its initial *μ* bases are sequenced they are used to identify its starting position and orientation along the reference genome, which determines the fate of the nascent fragment according to the current decision strategy (E). Upon rejection (upper path), an additional time of *ρ* sequenced bases later the pore is freed, a new read is acquired after further time *α*, and the model iterates from the beginning. Conversely, upon acceptance (lower path), the molecule translocates through the pore until all *l* of its nucleotides (i.e. *l* – *μ* additional bases) have been read. A new read can then be acquired and the model iterates from the start.

We call our novel method BOSS-RUNS, for “Benefit-Optimising Short-term Strategies for Read Until Nanopore Sequencing”. In brief, we quantify uncertainty at each site during a resequencing experiment, calculate the expected benefit of new reads, and dynamically adapt the sequencing effort to focus on fragments from areas of highest uncertainty. Our implementation efficiently communicates with the sequencing device through the readfish toolkit (Payne et al. 2021) and the Read Until API (Oxford Nanopore Technologies 2020) to incorporate the streamed sequencing data and provide updated decision strategies in real-time.

One promising application of our strategy is the possibility to compensate for the inherent tendency of some genomic regions to be sequenced at higher coverage than others, possibly due to GC content or other factors (Ross et al. 2013; Krishnakumar et al. 2018). In this scenario our method could lead to more homogeneous distribution of sequencing reads across a genome, with the benefit of increasing accuracy of genotype calling and reducing uncertainty in regions of low coverage. On the other hand, our method can also be used to focus the sequencing effort on ROIs or sites that show variation without the need for prior information about their location.

We demonstrate our novel method by mitigating coverage bias in a microbial mock community leading to higher coverage depth of low-abundance species, an increased limit of detection, as well as improved variant calling. To summarize, our approach of dynamic, adaptive sampling allows us to change what is sampled during an ongoing sequencing experiment as a result of the data that has already been collected and enables redistribution of coverage to sites of highest biological interest without *a priori* knowledge of coverage bias or variant sites.

## 2 Results

### 2.1 Quantifying information content of sequencing reads

In this work we present a framework that enables dynamic decisions during sequencing using nanopores. By calling it *dynamic*, adaptive sampling we describe our extension compared to current approaches, which are limited to *a priori* choice of target regions. In this section we give an overview of the decision framework, with further details and formal explanations provided in the Supplement (Suppl. Sect. 1.1).

First, we capture the amount of information we have at each site of a genome under investigation by considering a probability distribution over all possible genotypes of that site. We then update this probability distribution as we collect data throughout the experiment, i.e. we calculate a posterior distribution that includes the observed nucleotides from reads that cover that position. Next, we calculate the remaining uncertainty and how much information we might gain from one further sequencing read covering that site. By combining these scores over adjacent sites we can quantify the expected information gain from a sequencing read solely by knowing its starting location and its orientation. Finally, by ranking the expected benefit of reads and taking into account the expected time of sequencing them we can calculate the optimal subset of sites to accepts reads from in order to increase the gain of benefit at that moment in the experiment. The following paragraphs give more details on the individual steps involved to arrive at such a dynamic decision strategy.

#### Probability distribution of genotypes at each site of a genome

We start by defining a probability distribution of possible genotypes at each position of one or multiple genomes. Briefly, the genotype probability distribution takes both prior information about the genotype, e.g. from a reference genome, and already observed bases at a position into account. (Throughout, we use reference genome to describe any assembly used for a resequencing experiment and not necessarily a reference assembly that is representative of the investigated species.) Additionally, ploidy and probabilities of sequencing errors are considered.

Given already observed read data *D*, containing *n* reads covering position *i*, we denote by *d_j,i_* ∈ *B*, with *B* = {A, C, G, T}, the nucleotide in read *j* that maps to *i*. For a haploid genome, the set of possible genotypes is *G* = *B*, whereas for diploid genomes *G* instead consists of unordered pairs *g* = {*b*_1_, *b*_2_}, with *b*_1_,*b*_2_ ∈ *B*. For simplicity we present the case of genetic diversity and sequencing errors only occurring as SNPs; an extension that includes deletions and is used in our applications is provided in the Supplement (Suppl. Sect. 1.2). We define prior probabilities for genotype *g* at position *i* as *π_i_*(*g*), and the probability of calling base *d_j,i_* assuming genotype *g* as *ϕ*(*d_j,i_*|*g*), which represents a matrix of observation probabilities given assumptions about ploidy and sequencing errors (details in Suppl. Sect. 1.1). The posterior probability of genotype *g* ∈ *G* at *i*, conditional on *D*, is then

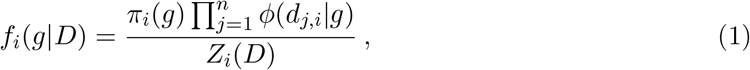

where *Z_i_*(*D*) represents a normalising constant, i.e. the likelihood of the data, that ensures the posterior probabilities sum to 1.

This model allows us to quantify the uncertainty about the genotype at each site (Fig. 1C) and in turn makes it possible to calculate the expected reduction in uncertainty resulting from observing a newly sequenced read. We call this expected reduction of uncertainty the *positional benefit score* of a site. This quantity summarizes the expected change in the genotype probability distribution given one additional observation at that position and is calculated as follows: given the current data (*D*) we imagine that we observe one additional nucleotide *n* at position *i*, i.e. *d*_*n*+1,*i*_, calling this augmented data *D′*. We then measure the difference between the distribution of genotype probabilities resulting from *D* and *D*′ by the Kullback-Leibler divergence (*D_KL_*; Kullback and Leibler 1951). Lastly, we sum over the different possible nucleotides *d*_*n*+1,*i*_, weighting their contributions by the estimated probability of observing them in the next read, to compute the expected reduction in uncertainty:

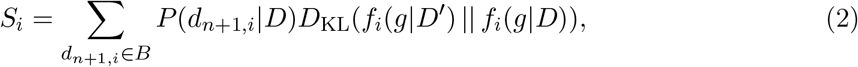

where the estimated probability of observing nucleotide *d*_*n*+1,*i*_ in the next read is given by

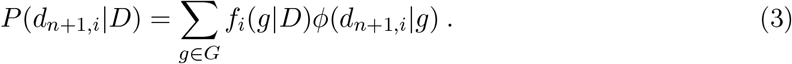

A practical way of calculating the positional benefit scores and some examples at different coverage patterns are given in the supplementary material (Suppl. Sect. 1.3, Suppl. Fig. 1). Broadly speaking, positions that are already covered by many agreeing reads will receive a low score and, conversely, positions covered by few, or possibly contradictory reads will score highly as individual observations have higher potential of influencing the probability distribution. This technique of defining the information gain in terms of the Kullback-Leibler divergence of two distributions is used in Bayesian experimental design (Chaloner and Verdinelli 1995) and is equivalent to evaluating the expected reduction in Shannon entropy (Shannon 1948) brought by a new read.

#### Estimating the expected benefit of sequencing reads

Since we want to quantify not only the remaining uncertainty at individual sites but also the potential information gain of future sequencing reads, we consider the fact that reads are derived from contiguous sections of a genome. So, as a next step, we combine the positional benefit scores across sites that a sequencing read might span, in order to evaluate the expected benefit of such a read (Fig. 1D). We assume that a sequenced read will cover a number of consecutive sites of a reference genome equal to the molecule’s length *l*. The expected benefit is then calculated as the sum of consecutive positional scores, beginning from the read’s mapping starting position *i*, weighted by the distribution of previously observed read lengths, *L*(*l*). In other words, we form the sum 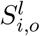 of consecutive positional benefit scores of a read of length *l* starting at position *i* with orientation *o* (*o* = 1 indicating a read in the forward direction relative to the reference genome, and 0 indicating the reverse direction); and then combine these, weighted by the probability that the read will reach that position (Fig. 1D). For a forward-oriented read 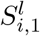 will be:

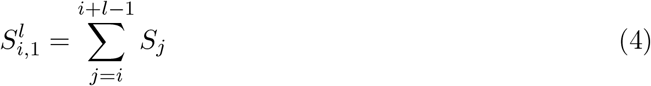

(see Suppl. Sect. 1.4 for the reverse-oriented case), leading to the expected benefit:

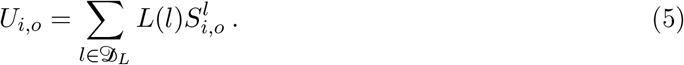

Here, 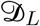 represents the domain of *L*(*l*), i.e. all read lengths observed so far. In reality we use a truncated normal distribution as a prior for read lengths, which we continuously update with observed lengths of full-length sequencing reads throughout an experiment. More details about *S^l^* and *L*(*l*) are given in Suppl. Sect. 1.4.

With this, we can quantify the expected information gain of a sequencing read solely on the basis of its genomic origin and orientation. Ultimately, a sequencing read that is expected to give a higher sum of scores, i.e. a greater reduction in the uncertainty of genotypes at the positions it covers, will be considered more useful than a read with a limited potential to alter the previously defined posterior probabilities. We provide an approximation to calculate this quantity efficiently based on a piece-wise approximation of the read length distribution in the Supplement (Suppl. Sect. 1.4).

### 2.2 Decision framework to enable dynamic, adaptive sampling

Using the expected benefit of reads we can now define a framework for making decisions about which fragments to sequence fully and which to reject from nanopores. Note that in line with common usage we refer to fragments, i.e. DNA molecules, and reads, their translation into sequence space, interchangeably. Overall, our aim is to optimise the rate of accumulation of information, i.e. of expected benefit, across all pores and over time. As we collect data throughout the sequencing experiment the value of reads at different positions will change, and therefore the decision strategy will have to adapt to these changes dynamically in real-time. Such strategies are stored as Boolean vectors and indicate the decision to be made about a sequencing read starting at any genomic position (Fig. 1E).

To define our decision strategies we parameterize the duration of individual steps in the sequencing process. As our time unit we use the amount of time it takes one base to translocate through a pore (Fig. 1F). Analogous to Read Until and readfish, we start sequencing a DNA fragment and use *μ* initial bases to determine its genomic origin and orientation. The value of *μ* is assumed constant in our model and can be adjusted to ensure mappings of sufficient quality, e.g. depending on the complexity or repeat content of the used reference genome. In reality *μ* depends on the size of individual data chunks used for real-time base-calling. The smallest useful setting is 0.4 seconds of input data, which corresponds to ~180 nucleotides assuming a translocation speed of 450nt/s. In our applications we used 0.8s of data and observed a mean length of 348nt for real-time base-called data chunks used to determine the origin of fragments. We further assume some constant time is needed to effect the rejection of a read (*ρ*) and to acquire a new read at a pore (*α*). In line with measurements from sequencing experiments our model assumes *ρ* = 300 and *α* = 300 by default. The effects of real-life conditions and constraints on these parameters are discussed in the Discussion. If a fragment is sequenced fully, time equal to its length *l* passes and benefit 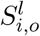 is accrued (with expectations *λ* = E[*L*] and *U_i,o_*, respectively); by rejecting a read, time equal to *l* – *μ* – *ρ* can be saved and the expected gain of benefit is limited to the position scores of its initial fragment, i.e. 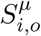 (see Fig. 1F and Suppl. Fig. 2).

#### Finding an optimal decision strategy to maximise information gain

With this parameterization of the sequencing process we determine an optimal sequencing strategy that maximises the expected benefit per unit of sequencing time given the currently available data. Such a strategy, denoted as 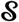, can also be seen as an indicator function that returns 0 or 1 for all combinations of genomic position and fragment orientation, e.g. 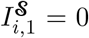 indicates the rejection of a forward-oriented read at position *i* and 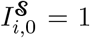 the acceptance of a reverse-oriented read. Our aim is therefore to find an optimal strategy 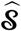 that maximises the benefit per unit time 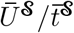 given the current data *D*:

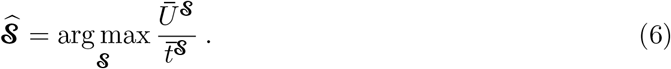

Here 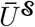 is the average expected benefit. Given a genome with a total length *N* and the average expected benefit of the initial parts of reads, i.e. the benefit accrued from the initial fragment used in the decision process, denoted 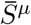, it takes the form:

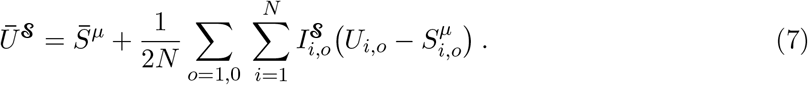

In other words, it is the sum of the average expected benefit from a read of *μ* bases and the average of a fully sequenced read, which adds further benefit of 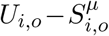 if the indicator function for that position-orientation combination returns 1. Then, 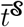 is the expected time needed to complete the processing (whether accepted or rejected) of a read:

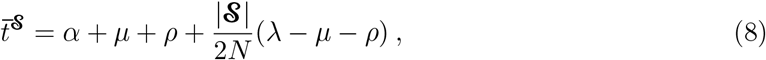

where 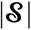 denotes the size of the strategy, i.e. the number of position-orientation pairs for which the indicator function will return 1, and λ is the mean read length (E[*L*], as above).

For simplicity, here we assume uniformity of the distribution of read origins; we present a generalisation used in our implementation in the Supplement (Suppl. Sects. 1.5 and 1.6).

To compute the optimal strategy, we rank all of the position-orientation combinations (*i*, *o*) in decreasing order of the expected benefit gain from sequencing them in their entirety 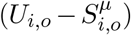. Starting with an empty strategy (one that rejects all reads) we successively include the ranked sites and test after each one whether its contribution results in an improvement over the previous strategy, i.e. whether the current iteration achieves higher gain of benefit per time unit 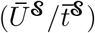 than the preceding strategy that included one fewer site (position-orientation pair).

We call our approach of finding an optimal strategy BOSS-RUNS: “Benefit-Optimising Shortterm Strategy for Read Until Nanopore Sequencing”. For further details, an overview of parameters and variables in the model and proof of optimality please see Suppl. Sects. 1.5 and 1.7 and Suppl. Table 1.

### 2.3 Real-time implementation

Our novel method is implemented to interact with a version of readfish (Payne et al. 2021) modified to read updated target lists throughout the experiment (available from https://github.com/LooseLab/readfish/tree/BossRuns/V0.0.1). The sequencing device performs real-time basecalling and deposits fastq files containing batches of reads (4000 by default). BOSS-RUNS monitors the device’s output and periodically (by default every 30 seconds) includes all new data in an updated decision strategy. In this process new coverage counts are incorporated after mapping basecalled reads to one or more reference genomes using mappy, the python wrapper of minimap2 (Li 2018). If a read maps to more than one position, the best alignment is chosen based on mapping quality or the alignment score of the dynamic programming algorithm in case of a tie. Observed lengths of fully sequenced reads and their mapping positions are continuously used to update the empirical distributions of read lengths *L*(*l*) and read start locations and orientations *F_i,o_*. To prevent the strategy from getting too greedy updates are only applied when a region surpasses a threshold of average coverage (default: ≥5× in 20kb windows).

In order to keep up with the real-time data stream and to ensure optimality of the strategy at any point in time, new results need to be calculated quickly. For this we use several optimisations including a fast algorithm to find approximate decision strategies, described in Suppl. Sect. 1.8. Our method can use either single or multiple reference chromosomes/genomes as input and optional masks to indicate initial ROIs, similarly to current approaches to adaptive sampling (Payne et al. 2021; Martin et al. 2022). In that case the scope of the dynamically updated strategies is limited to the ROIs and flanking regions around them and reads originating outside will always be rejected. If multiple references are considered, the expected benefit of reads is calculated separately per reference and then used to derive a common decision strategy across all considered references. This is to ensure that we can account for differences in the distributions of read lengths and read starting positions between genomes, while also sequencing the most informative reads of a mixture instead of focusing on the most informative reads of each individual genome or chromosome.

BOSS-RUNS is implemented in python and available at https://github.com/goldman-gp-ebi/BOSS-RUNS. We provide a conda environment for its dependencies: readfish (Payne et al. 2021), ONT’s MinKnow API (4.2.4; Oxford Nanopore Technologies 2021), numpy (1.21.1; Harris et al. 2020), numba (0.53.1; Lam et al. 2015), scipy (1.7.1; Virtanen et al. 2020), mappy (2.22; Li 2018), pandas (1.3.3; McKinney 2010), toml (0.10.2; Pearson 2022), and natsort (7.1.1; Morton 2021).

### 2.4 Dynamic enrichment of differentially abundant species

#### Experimental setup

Enrichment of ROIs by rejecting unwanted reads has been previously demonstrated (Loose et al. 2016; Kovaka et al. 2021; Payne et al. 2021). BOSS-RUNS can be applied more generally, and makes use of targeted rejections even in the absence of specific ROIs. Here we consider a scenario of whole-genome resequencing where the entire genome is considered of interest, while we showcase a scenario with ROIs in the Supplement (Suppl. Sect. 2.1).

One situation where the possibility of redistributing data is very effective is in the presence of coverage bias. This can be the case when sequencing either a single organism or multiple genomes. Our first experiment had two major goals. Firstly, we aimed to mitigate bias in coverage across multiple differentially-abundant genomes. Secondly, we demonstrate that using a dynamic approach to adaptive sampling can increase the sampling of sequencing reads from variant or difficult-to-resolve sites without prior knowledge of their location. We therefore sought to sequence eight bacterial species of the well-characterised zymoBIOMICS microbial mixture (ZymoBIOMICS DNA Standard II D6311, Zymo Research). The abundance of the organisms in this mixture is logarithmically distributed with the most abundant species, *Listeria monocytogenes*, comprising 90% of total DNA, the second most abundant *P. aeruginosa* circa 9%, the third (*B. subtilis*) 1%, etc. (Fig. 2A). To measure the effects of BOSS-RUNS we divided the available pores on the flowcell and used our new method on one half while the remaining pores did not perform any read rejections and therefore acted as control.

**Figure 2:**
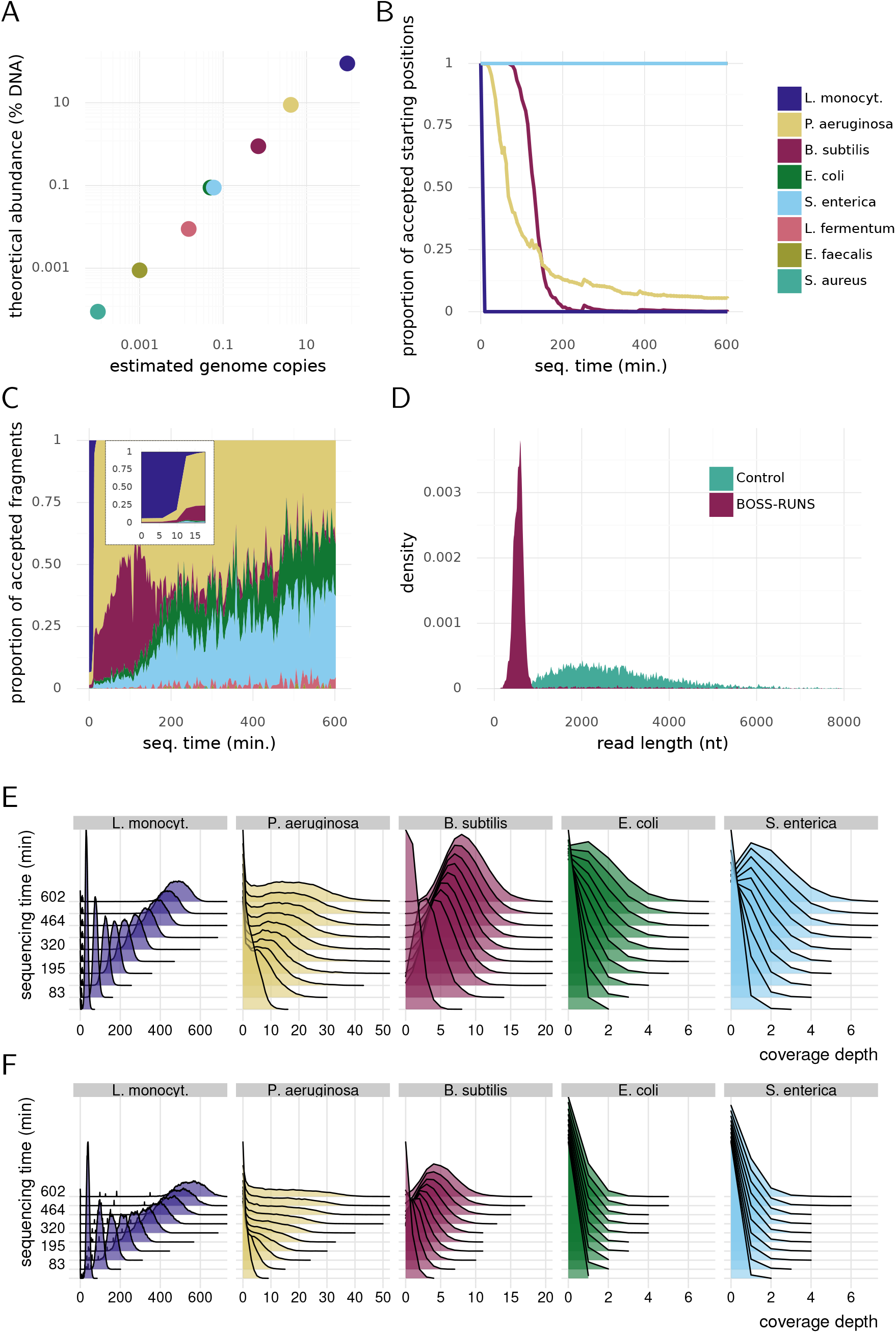
BOSS-RUNS can dynamically adapt to the differential abundance of bacteria in a mixture. A) We sequenced eight bacterial species of the zymoBIOMICS mixture with logarithmically distributed abundances covering seven orders of magnitude. *E. coli* and *S. enterica* are equally abundant at 0.1% of total DNA in this microbial community. Colors correspond to species as in (B–C) and (E–F). B) Starting with a strategy that accepts reads from any position in all considered genomes, we quickly observe rejections of reads from the most abundant bacteria *L. monocytogenes*, followed by *P. aeruginosa* and *B. subtilis*. The plot shows the proportion of position in each species’ genome for which a read starting there is accepted by the current BOSS-RUNS strategy, over the duration of the experiment. C) The proportion of the accepted fragments that derive from each of the different bacterial strains demonstrates the effect of the changing decision strategy in (B). The inset plot, focusing on the start of the experiment, shows how the strategy rejects almost all *L. monocytogenes* reads after the first 10min. D) The distribution of read lengths confirms that BOSS-RUNS rejects the majority of sequencing reads, with a clear peak that corresponds to the initial part of a sequencing read used in the decision process. E and F) The coverage distribution across all species using BOSS-RUNS (E) shows depletion of DNA from more abundant genomes in turn for enrichment of information from rare species, compared to the control section on the flowcell (F). Plots show the distribution of coverage depths over sites, with different layers within a plot indicating different time points within the experiment. Accumulation of coverage over time is shown by the distributions’ shift to the right within each panel; note the decreased, but still very high, coverage of *L. monocytogenes* using BOSS-RUNS, and the increased coverage for the other (less abundant) species. Results from the three least abundant bacterial species are omitted due to the differences not being obvious in this type of visualisation.

To mimic a realistic sequencing experiment where we do not have prior knowledge about the exact bacterial strains contained in the mixture we used assemblies of closely-related strains instead of accurate assemblies generated from the strains known to be present in the community (see Sect. 4.2 for accessions). Additionally, this allowed us to evaluate the performance of our method in focusing on sites that differed between the reference genomes we used and the true genomes. Priors of genotype probabilities were also initialised using these references.

#### BOSS-RUNS’ strategy

During the sequencing experiment we can observe how the decision strategy changes over time. As the genomes of individual bacteria are continuously resolved, i.e. we become more certain about the genotype at many sites, the proportion of positions at which we still require more information decreases. Due to the differential abundance of the considered species, *L. monocytogenes* is considered mostly resolved after only a few minutes followed later by *P. aeruginosa* and *B. subtilis* (Fig. 2B). Accordingly, the proportion of accepted reads demonstrates that the focus switches from the most abundant bacteria towards rarer species (Fig. 2C). Interestingly, the rate at which individual genomes are resolved is not equal across all bacteria in the mixture. For example, the proportion of fragment start sites from which reads would be accepted from *L. monocytogenes* or *B. subtilis* decreases to values close to 0. On the other hand, *P. aeruginosa* approaches a level of ~5.8% at the end of the experiment and does so at a slower rate. In other words, some sites in the genome of *P. aeruginosa* require more data to be confidently resolved and a portion of sites remains uncertain despite sampling data throughout the run. This is in part due to different levels of divergence between the strains in the zymo community and the reference genomes we used during the experiment, and in part due to differential effects of coverage bias also within each species’ genome.

Given the large difference in abundance and the prompt resolution of *L. monocytogenes*, we expect the majority of sequencing reads to be rejected throughout the experiment. Indeed, the distribution of observed read lengths confirms that BOSS-RUNS ejects most molecules resulting in a peak at ~480bp (Fig. 2D). The bimodality of the read length distribution after separating the sequencing data by target species also corresponds to our expectations given the proportion of rejected reads from each species (Suppl. Fig. 3). Interestingly, the presence of a peak at around 480bp in the read length distribution of rare species indicates that some reads from these species are also rejected. The majority of these false rejections (84%) was due to inability to determine the source species from the initial fragment. Notably, sequencing reads in this experiment were generally relatively short due to the necessity of preceding amplification.

#### Improved sequencing of bacterial species

The effect of the changing decision strategy becomes evident when looking at the distribution of coverage depth over time. Sequencing coverage from the most abundant species is effectively redistributed to the other scarcer species compared to the control (Fig. 2E, F). For example, for *E. coli* and *S. enterica*, which comprise only 0.1% of the input DNA, we achieve 3.9 and 4 times the coverage compared to the control.

Changes in mean coverage over time confirm these observations regarding the overall shape of the coverage distributions. Indeed, sacrificing data from heavily sampled organisms enables us to obtain more DNA from rare species (Fig. 3A). For example, BOSS-RUNS achieves between 4.1 and 5.8 times more coverage of the scarce bacteria (*E. coli*, *S. enterica*, *L. fermentum*, *E. faecalis*). Focusing on low-coverage sites rather than mean coverage, the proportion of sites with coverage less than 5 × also highlights the advantage gained by using our dynamic decision strategies. This quantity decreases quicker, and reaches lower final levels, compared to the control for all but the most abundant genome (Fig. 3B). The redistribution of sampled sequencing data from regions already well-covered to areas of low coverage is one of the main features of BOSSRUNS. In the case of *B. subtilis*, for example, this leads to less than 5% of sites with coverage less than 5× with BOSS-RUNS, against ~44% for the control. In rare species we checked whether the sites with a coverage of more than 5× were caused by reads mapping to repeats or other regions of low complexity. Analysing the overlap of these sites and the respective repeat-masked genomes revealed that this was not the case and the observed coverage at these sites was not artefactual but true enrichment (Suppl. Table 2).

**Figure 3:**
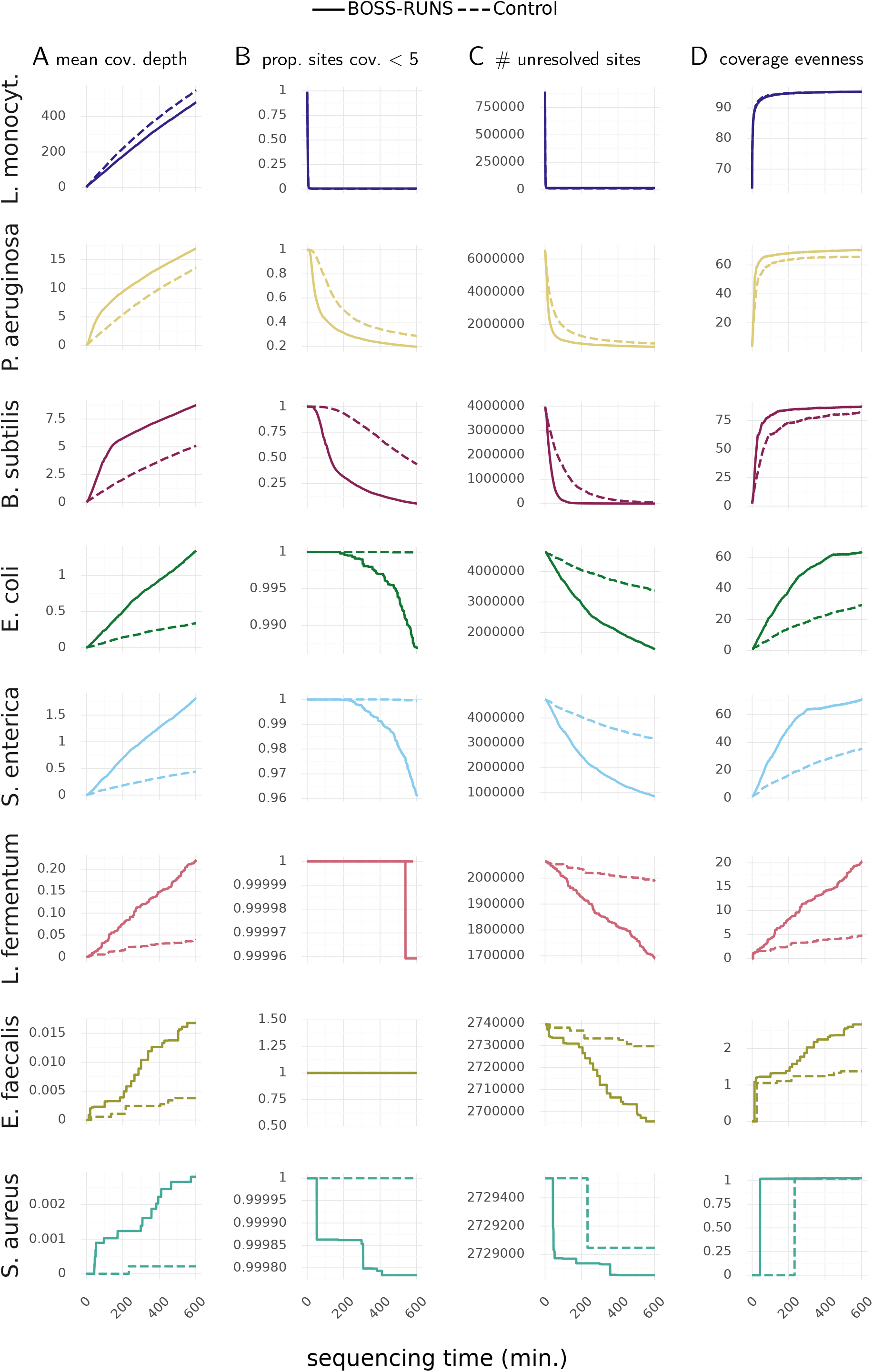
Various metrics demonstrate the improvements in sequencing the bacterial community using BOSS-RUNS (solid lines) and control (dashed lines) on the same flowcell in an experiment lasting 10h. A) The mean coverage depth across the genomes of all considered species. Sequencing coverage of more abundant species will generally be traded-off by BOSS-RUNS to collect more data from rare species. Upon (mostly) resolving individual genomes a change in the rate of data accumulation is visible, e.g. after ~180min for *B. subtilis*. B) The proportion of sites that remain at <5× depth reveals that data is efficiently redistributed by BOSS-RUNS to areas of low coverage. C) Using the genotype posterior probability distribution, we can classify sites as resolved if the probability of one genotype is >0.99. Across all species BOSS-RUNS achieves lower numbers of unresolved sites throughout the experiment owing to both sampling more data from rarer species and redistributing data within each genome. D) Coverage evenness describes the homogeneity of observed data across a genome independent of absolute coverage. By focusing the sequencing effort on sites with low coverage our method leads to more even distribution of coverage. Note the different scales on the *y*-axes of all plots, to allow for the sampling statistics of species of widely varying abundances.

We can also use the uncertainty about each sites’ genotype, as defined in our model, to classify sites into two groups, unresolved and resolved. When the posterior probability of one genotype at a site surpasses 0.99 we declare that site as resolved, and can count the sites which still require more data to reach that level of certainty. Again BOSS-RUNS shows higher performance by reaching lower numbers of unresolved sites in a shorter time (Fig. 3C).

Balancing coverage bias across genomes is not the only benefit of our new method though, as data is also redistributed within individual genomes. This effect is partly responsible for the gains described so far, but is overshadowed by the stark species abundance differences. By using a measure of evenness that describes the uniformity of coverage distribution and is relatively independent of the absolute coverage (Mokry et al. 2010), we observe that BOSS-RUNS not only boosts the coverage of rare species, but also ensures that coverage is more uniform in all species, including those of higher abundance (e.g. *P. aeruginosa* and *B. subtilis*, Fig. 3D). Even in cases where the total collected coverage of a bacterial strain might be lower it is possible that this more uniform distribution of coverage and focus on sites with higher uncertainty could achieve a more desirable outcome of the experiment with BOSS-RUNS, although in our experiment this effect is not readily visible in Fig. 3 for *L. monocytogenes*, we note that the improved precision of SNP detection for this species (see below; Fig. 5B) could be due to these effects.

#### Redistributing coverage to under-sampled sites

Another way to explore the redistribution of data within genomes is to examine the already observed coverage at the sites that a read maps to when the decision about that read was made. If our method indeed successfully focuses on reads from areas with highest uncertainty we expect the mean coverage at sites spanned by accepted reads to be lower than sites spanned by rejected reads. An even bigger effect might be visible when looking at the minimum coverage within those spans, since the decision to accept reads might be driven by low coverage at individual sites as opposed to low average coverage in an area. To investigate this, we separated the reads by the decision made during the experiment: either to sequence them entirely or to reject them. We then record the mean and minimum coverage of all sites that a read maps to and pool those measurements at different timepoints.

To demonstrate BOSS-RUNS’ effects, we focus on the results for a single species, *B. subtilis*, for which the decision strategy accepts all reads until ~80min into the sequencing experiment; then rejects an increasing proportion of reads; and is rejecting most, but not all, reads from ~200min onwards (see Fig. 2B). Looking at the background mean coverage at sites spanned by accepted and rejected reads, we see an overlap up until the size of the strategy starts to diminish (Fig. 4, left). Throughout the rest of the experiment, as expected, both the mean and minimum coverage of sites spanned by accepted reads are consistently lower than for rejected reads, albeit with larger variation due to the decreasing number of fragments sequenced in their entirety (Fig. 4). In summary, this shows that despite this genome being mostly resolved early in the experiment, we continue to sample fragments covering positions of low average coverage or where we only have minimal information about individual sites. These results confirm that BOSS-RUNS focuses on these reads not solely due to the abundance difference but also due to the coverage variation within the data of *B. subtilis*. Notably, it might be surprising that we observe a few rejections of reads originating from this species right from the start of the experiment. This is due to reads failing to map when using only their initial parts during the decision process.

**Figure 4:**
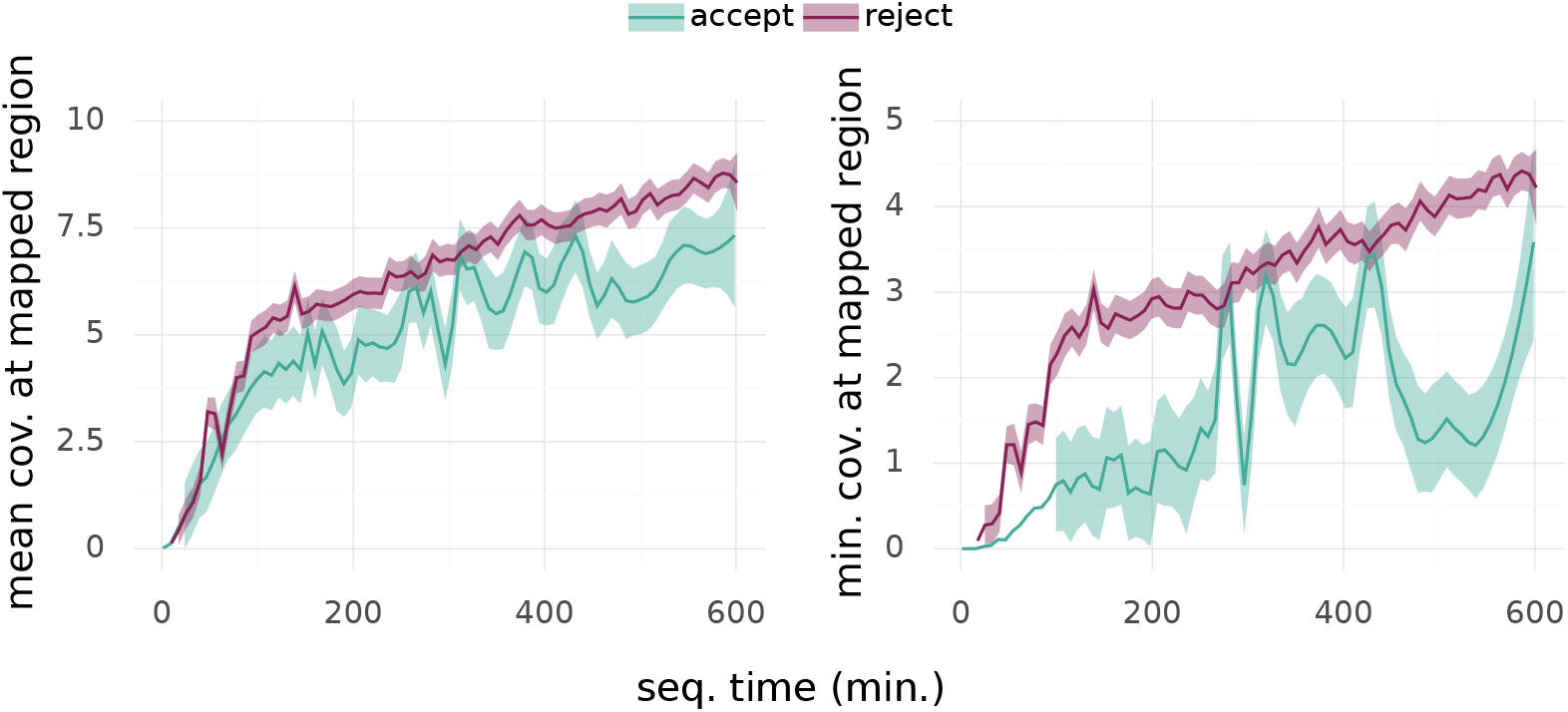
Coverage is effectively redistributed by sampling more data from sites with low coverage. Focusing on *B. subtilis*, the mean observed coverage of reads’ mapping locations first overlaps for rejected and accepted reads (left). As more fragments are rejected from pores, the mean coverage of sites spanned by accepted reads becomes, and stays, consistently lower than for rejected ones. We observe an even larger difference in the minimum coverage of regions reads map to (right). In combination, these demonstrates that data is continually sampled from areas of low coverage even after most of this species’ genome has already been resolved.

### 2.5 Focused sequencing leads to improved variant calls

Next, we sought to perform variant calling for five of the eight bacterial species in the microbial mixture. (We excluded the three least abundant species from this analysis, as we did not collect enough data to make reliable calls.) With this analysis we tried to answer the questions of (a) whether we could successfully sample data from rare species in order to better identify the differences between reference assemblies and the true sample genomes, and (b) whether BOSS-RUNS can effectively focus on sites where we observe variation and therefore increased uncertainty.

Our analysis is based on comparing inferred variants from data accumulated using BOSSRUNS (or the control) to a ground truth derived from deep, short read sequencing of the same bacterial strains (see Methods, Sect. 4.3). By making comparisons at multiple timepoints throughout our experiment, we show how knowledge of variants accumulates over time (Fig. 5); in future, such information could be used to shorten the duration of experiments needed to achieve particular levels of accuracy. For the most abundant species, *L. monocytogenes*, the decreased coverage with BOSS-RUNS leads to slightly lower sensitivity than for the control case for that species. Nevertheless, high sensitivity is achieved in a very short time, and the effective redistribution of coverage within this species’ genome leads to increased precision. In turn, however, for all the other species the increased and better-targeted coverage produced by BOSS-RUNS (see above) means more variants are discovered, with improved sensitivity and precision.

**Figure 5:**
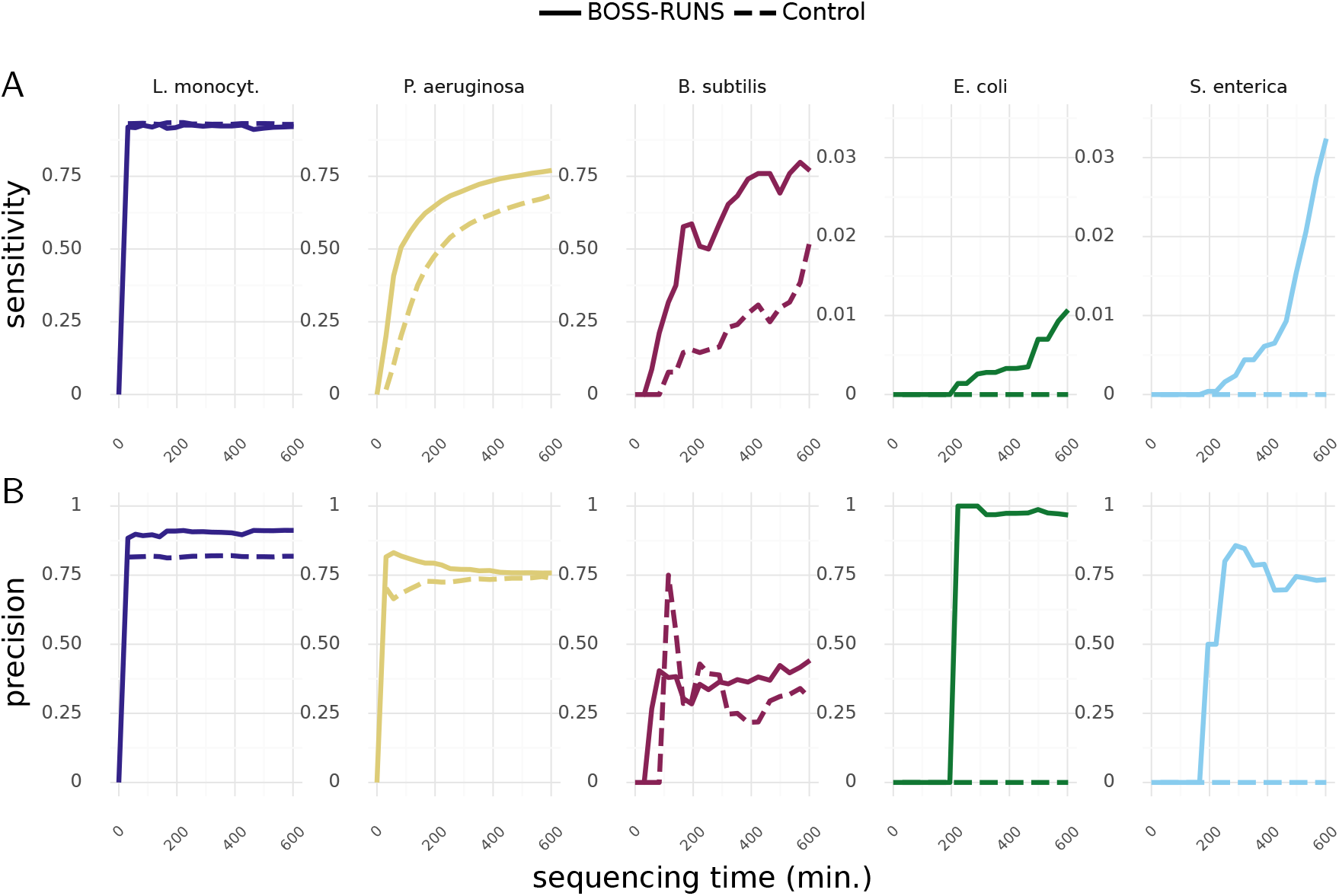
Dynamic, adaptive sampling leads to improved discovery of SNPs. We compared the variants called from data collected without adaptive sampling (control) and with BOSS-RUNS to a set of ground truth variants from deep, short read sequencing of the same bacterial strains. By performing variant discovery at different timepoints we can gain further insight into the advantages of our method. A) Whereas the sensitivity of BOSS-RUNS is slightly lower for the most abundant species *L. monocytogenes*, we observe a larger number of discovered true positives in all remaining genomes. To highlight differences, we set the y-axis ranges to 0–0.95 for the first three species, and 0–0.035 for the remaining two. B) The precision of variants called from data generated using BOSS-RUNS is at a similar level to the control or moderately higher.

Interestingly, even for the two bacteria *P. aeruginosa* and *B. subtilis*, which are considered mostly resolved by our method during the experiment leading to the majority of reads being rejected, we still see an increase in sensitivity at later stages of the run (Fig. 5A). This is due to the ability of our method to sample more data specifically at positions where it is conducive to reducing uncertainty about the genotype. For example, after 10h of sequencing BOSS-RUNS finds 26,481 variant sites in *P. aeruginosa* (sensitivity 0.79), whereas we observe 23,541 SNPs from control data (sensitivity 0.68), despite the decision strategy rejecting fragments from >80% of the genome after the first 3h. At the same time the precision of variant calls on the data collected with BOSS-RUNS is either moderately higher or at a similar level to the control dataset (Fig. 5B). In the rarer species the advantage of BOSS-RUNS simply collecting more data is evident as we are able to call SNPs at least in some regions (119 and 80 SNPs detected after 10h for *E. coli* and *S. enterica*, respectively), whereas the control data does not contain enough reads to produce any variant calls.

## 3 Discussion

In this work we have shown that our novel approach to dynamic, adaptive sampling for nanopore sequencing, implemented in BOSS-RUNS, provides a mathematical framework and fast algorithms to generate decisions strategies that optimise the rate of information gain during resequencing experiments. Compared to sequencing without adaptive sampling, this leads to an increase in the sequencing yield of on-target regions, specifically at positions of highest uncertainty. Additionally, it can effectively mitigate abundance bias or other sources of non-uniform coverage, for example from enrichment library preparation procedures, leading to smaller proportions of sites at low coverage depth and greater evenness of coverage. Furthermore, our methods can lead to improved discovery of variants by both sampling more data from negatively biased regions or species in the input material and by focusing the sequencing on sites where the underlying genotype is not clear from the data observed up to that point in time. Overall, our method can help to shorten the time-to-answer and reduce the cost of sequencing.

Current methods of adaptive sampling base the rejection of sequencing reads solely on whether they cover some ROIs that were defined before starting the sequencing run. Instead, we can change our targets throughout an experiment in order to collect data where it is most useful to reduce the uncertainty about the underlying genotypes at each site.

We demonstrated the effectiveness of our approach by sequencing a microbial mixture affected by coverage bias. We show that BOSS-RUNS adapts to the differential abundance of the targeted genomes without prior knowledge about the composition of the input library. It not only reduces the inter-species bias, but also redistributes coverage effectively within-species.

In common with any resequencing experiment, the only piece of prior knowledge we do require is a reference genome that is related to the organism(s) we expect to observe in the sequenced material. This is used for two purposes. Firstly, we initialise the genotype prior probabilities according to the reference. These can either be informed by the nucleotides present in the chosen reference or they can be uniform for all possible genotypes, perhaps depending on how much divergence we expect in the input material. Priors signifying different amount of confidence in the chosen reference could also be used, but are not implemented due to the negligible effect of the priors in many cases (see Suppl. Fig. 1). Secondly, the reference genome is used to determine the origin and orientation of nascent sequencing reads. This is the basis for making decision: barring major structural rearrangements, moderate amounts of divergence will not have a substantial impact on the performance of BOSS-RUNS.

Determining the mapping location of the initial part of a read using minimap2 does not achieve perfect sensitivity or precision, and indeed we see some reads rejected due to them not mapping with sufficient confidence (see Suppl. Fig. 3). Admittedly, using minimap2’s preset parameters for long reads is not ideal in this case and optimising parameters to better accommodate such rather short reads but with the error profile of long reads could reduce the number of falsely determined off-target reads. Alternatively, other methods for read classification could be used such as the recently described ReadBouncer, based on the DREAM index (Ulrich et al. 2022).

However, our combination with readfish, which uses GPU-accelerated high-accuracy basecalling, seems to be affected to a lesser degree than ONT’s built-in adaptive sampling, which has been reported to falsely reject more than one third of on-target reads in some cases (Martin et al. 2022).

In our experiment sequencing a mock bacterial community we use reference assemblies that are closely related to the actual strains contained in the mixture but not identical, to mimic a real setting where we do not know the true composition in advance. We measured their divergence in terms of the percentage of aligning nucleotides and ANI values using JSpecies (Richter et al. 2016), which range from 86.07 to 99.7% and 98.82 to 99.92, respectively (Suppl. Fig. 4). We can see that the divergence between the chosen reference and the true experimental strain has potential implications on the performance of BOSS-RUNS. For example, the reference we used for *P. aeruginosa* harbours more differences from the true genome than there are between the reference and true genome of *L. monocytogenes*, leading to the visibly different respective plateaus of remaining unresolved sites approached during the experiment (Fig. 3C). An important influence in the case of prokaryotes is the concept of pangenomes and the possibility of differential composition of accessory genes (Medini et al. 2005; Ozer et al. 2014). If a mixture contains more than one strain of a bacterial species we expect that sequences from all strains will be enriched, so long as they are contained in the chosen reference, as we will not be able to differentiate their origin with enough resolution. Indeed, Martin et al. (2022) have shown that targeting a single strain of *E. coli* also led to the enrichment of four other strains contained in the same gut microbiome standard. This phenomenon might also limit the advantage of BOSS-RUNS in extreme cases, since we will not be able to resolve regions of an assembly that are missing in the input library. We have implemented ways for BOSS-RUNS to ignore such missing areas, but still need to improve these techniques in order to better differentiate between absent parts of the genome and very low or spurious coverage, e.g. to prevent continuously waiting for reads from small parts of a genome in the hopes of resolving them, as visible in Fig. 2B for *P. aeruginosa*. Alternatively we could also implement the use of pan-genome graphs to account for variable composition of strains instead of linear assemblies (Colquhoun et al. 2021). For eukaryotes this limitation will have a much smaller impact due to the generally larger proportion of core vs. accessory gene content (Brockhurst et al. 2019).

Another possible limitation of our method comes from the fact that strategies generated by BOSS-RUNS are optimised to gain the most benefit in the short term. Early on during a sequencing run they may reject fragments that would be considered relatively more useful later on, which might seem counter-intuitive and can make the computed strategies greedy (in the sense described by Jones and Pevzner 2004). As an example, we might reject reads from regions with average uncertainty at one point in time to focus instead on areas of high uncertainty; however, regions with average uncertainty could themselves turn into regions with relatively high uncertainty as the experiment continues: past rejection of reads from these areas might not have been advantageous in the long term. Further work will be needed to address this issue.

As mentioned above, the read lengths in our applications (modelled by the distribution *L* with mean λ) are relatively short with a mean of 3.11kb, which was in part due to the need for amplification in order to achieve sufficiently high molecular weight DNA. But this might serve as a proxy for the challenging nature of extracting DNA from metagenomic samples, which often relies on harsh, multi-step procedures to ensure that cells from all contained species are lysed and genomic material is available for sequencing (Quick 2019) and can lead to reads that are shorter than desired. For such short fragments the time needed for processing, rejection and acquisition of an alternative read (*μ* + *ρ* + *α* in our model) might approach the elapsed time for fully sequencing a typical read in its entirety. If reads were even shorter, using adaptive sampling might even be detrimental to the total amount of collected sequence. Martin et al. (2022) have recently modeled the influence of read length of the input library on the practically achievable enrichment (using Read Until but not dynamically updated strategies) of differentially abundant species in a mixture. They showed that average read length is a major determinant of the maximum level of enrichment and that the longer the reads, the larger the potential enrichment becomes (Martin et al. 2022). This may explain the moderate level of overall enrichment of the yield in our sequencing experiment.

Other factors that could influence the advantages of our approach include the availability of DNA molecules for sequencing at the flowcell surface. Whereas we did not observe differences in the acquisition time of new fragments at pores, we could imagine that experimental conditions might influence this parameter, for example DNA concentration throughout a sequencing run. In addition, the effects of variation of the duration and magnitude of the voltage reversal for effecting rejections have not been well-characterised. Exploring different experimental conditions and parameters in the context of adaptive sampling poses interesting questions for future studies of this technique.

A further consideration is any negative effect of repeatedly rejecting fragments on the status of nanopores on the flowcell. Pores might get blocked by DNA molecules that are not successfully ejected, which can reduce the number of active pores when using adaptive sampling (Loose et al. 2016). The use of nuclease flushed across the flowcell in order to free such clogged pores has been shown to mitigate this issue and recover blocked pores for sequencing (Payne et al. 2021; Martin et al. 2022). In our experiment, however, we did not observe any indications that the performance of the section on the flowcell running BOSS-RUNS was degrading faster than the control section when comparing the number of data-transmitting channels and their idling times (Suppl. Fig. 5, Suppl. Fig. 6) and others have reported similarly small effects on pore health (Martin et al. 2022). Depending on factors such as the proportion of rejected reads, read lengths and most importantly the total duration of the experiment, the impact on pore health might vary and especially for long sequencing experiments nuclease flushing could be important to retain a high number of active pores.

In the future we would like to apply our method to a wider array of biological problems. For example, we could adapt our model to quantify the uncertainty about the presence of epigenetic modifications, such an methylation, which are now becoming analyzable in real time with nanopore sequencing (Simpson et al. 2017; Leger et al. 2021; Liu et al. 2021; Zhang et al. 2021). In that case we could also inform priors with sequence context, in order to take into account that some modifications such as 5-methylcytosine (5mC) occur most frequently at CpG dinucleotides (Ehrlich et al. 1982; Bird 1986). BOSS-RUNS could also be used to overcome biases that are inevitably introduced by some library preparation methods. In whole-exome sequencing, an example of one of the most widely used resequencing techniques, differential pull-down efficiencies of genes and also different exons within genes can lead to substantial coverage bias (Barbitoff et al. 2020) which BOSS-RUNS could adapt to during an experiment and therefore help mitigate.

Further, we plan to extend our model to collect additional information besides the observed nucleotides at each site. For example, we could record the distribution of read lengths in different areas of the genome to focus sequencing effort on covering an entire genome with sufficient number of very long reads in order to maximise the contiguity of assembled contigs. Creating an approximate assembly in real-time could also eliminate the need for a reference genome when making decisions with BOSS-RUNS. We could also use linkage information from observing adjacent variants in the same sequencing read in order to express information about which haplotype the observed variation comes from in diploid or polyploid samples. BOSS-RUNS could then enrich for fragments that have the potential of bridging across variants and therefore help construct haplotype-resolved assemblies, at least in genomic regions with enough diversity. This could aid in the ongoing effort of obtaining completely phased *de novo* genomes from single individuals, and might reduce the complexity of currently used sequencing protocols or combinations of methods (Soifer et al. 2020; Porubsky et al. 2021).

Other innovations of nanopore sequencing in general might include the ability to repeatedly translocate the same native genomic fragment, which has been previously demonstrated for solid-state nanopores albeit without a nucleotide readout (Gershow and Golovchenko 2007), and more recently also by using helicases to translocate a DNA-peptide construct through protein nanopores (Brinkerhoff et al. 2021). If this was possible in a controlled fashion, the methods described here could be adapted to decide dynamically which reads to recapture and how many times to sequence them until their information content is exhausted and the sites they cover are resolved to some desired accuracy.

To conclude, our method expands the applicability of adaptive sampling and can potentially improve the information gain in many standard scenarios by dynamically enriching areas of highest uncertainty, e.g. genomic regions with increased divergence, discrepancy to expectations or simply lower coverage depth than the average. This is achieved by utilizing information derived in real time throughout a sequencing experiment, and can be used without prior knowledge of variation of the sample from a reference sequence. Ensuring more homogeneous coverage and overall less uncertainty about genotypes by focusing on biologically interesting sites leads to improved efficiency of sequencing using nanopores in a wide range of applications. The resulting reduction in the time-to-answer might be critical in a clinical setting or in pathogen surveillance.

## 4 Methods

### 4.1 Configuration of sequencing experiments

Sequencing was conducted on an ONT GridION using R9.4 flowcells. Since the quality and number of active nanopores can vary between flowcells it would be difficult to compare experiments involving adaptive sampling performed on multiple flowcells. Therefore we separated a single flowcell by assigning 256 channels to each of two different conditions. One of these two regions used a decision strategy that continuously accepts any encountered read, i.e. a control sector not performing any adaptive sampling, whereas the other was acting according to the decision strategies provided by BOSS-RUNS. Readfish was configured to reject reads from this sector if they did not map or mapped to (one or more) off-target sites, i.e. sites not included in the current decision strategy, or if no sequence was obtained from a fragment. For all our experiments we used 0.8s of data to infer the genomic origin and orientation of fragments before making decisions, i.e. roughly 350bp (corresponding to *μ* in our model, Fig. 1F), which results in a mean read length of 482bp for rejected reads due also to the additional time (*ρ*) taken to process and effect decisions. BOSS-RUNS deposits new strategies as compressed Boolean numpy arrays for each genome or chromosome, which are subsequently reloaded by readfish upon file modification. Communicating rejection signals to the sequencing device is performed by readfish.

### 4.2 Sequencing and analysis of the ZymoBIOMICS microbial reference

Input DNA from the ZymoBIOMICS Microbial Community DNA Standard II (Log Distribution D6311, Zymo Research) was prepared using SQK-LSK110 (ONT) and PCR amplified using the PCR expansion kit EXP-PCA001 (ONT). BOSS-RUNS and readfish depend on reference genomes to infer the origin of sequencing reads. In order to create a more realistic scenario where we do not know the exact bacterial strains, we decided not to use reference genomes from the strains contained in the microbial mixture, but instead used closely-related reference genomes identified by McIntyre et al. (2019). The employed assemblies are available in the European Nucleotide Archive under accessions ASM14656v1, ASM584v2, ASM400627v1, ASM39716v1, ASM30761v1, ASM51030v1, ASM25313v1, ASM810v1. Basecalling was performed using Guppy (5.0.16), set to high-accuracy mode.

To test whether increased coverage of rare species was due to repeats or low-complexity regions, we used RepeatMasker (4.1.2, default parameters; Smit et al. 2015).

### 4.3 Variant calling of bacterial species

To perform variant calling we used sequencing reads separated by their species of origin (using minimap2; Li 2018) and further partitioned them to comprise the cumulative data from the beginning of the experiment up to and including 20 individual timepoints, each separated by approximately 30min of sequencing (using custom python scripts).

To create a set of high-confidence variants we used publicly-available deep coverage short read sequencing of the zymoBIOMICS microbial community with evenly distributed abundances, which contains the same strains as the logarithmically distributed mixture (Zymo Research D6306). These data are available in the European Nucleotide Archive under the accession SRR13224035. Briefly, we mapped the separated reads to their respective assemblies (see previous section) using minimap2 (2.22; Li 2018) and samtools (1.12; Danecek et al. 2021), marked duplicates using picard (2.26.6, default parameters; Broad Institute 2019), and called variants, i.e. the differences between the assemblies we used and the strains contained in the sequenced microbial community, with freebayes (1.3.5, default parameters; Garrison and Marth 2012). Variants were filtered by minimum depth of coverage of 20 and quality score 20, transformed into their primitive constituents (vcflib, 1.0.2; Garrison et al. 2021), and sorted using bcftools. Variant calling from nanopore data of the zymo microbial mixture was done using medaka (1.4.3, default parameters, model r941_prom_hac_variant_g507; Oxford Nanopore Technologies 2022). For subsequent comparisons of vcf files we used vcfeval (rtg-tools, 3.12.1; Cleary et al. 2015).

## Data access

The source code of BOSS-RUNS is available at https://github.com/goldman-gp-ebi/BOSS-RUNS. The sequencing data generated in this study have been submitted to the ENA database under accession number PRJEB51967.

## Acknowledgements

We would like to thank Alex Payne for valuable insights and helpful discussions. This work was supported by the Biotechnology and Biological Sciences Research Council [grant number BB/N017099/1 to ML]. LW, NDM, CM, EB and NG were supported by the European Molecular Biology Laboratory. CM was also supported by Murray Edwards College, Cambridge, and by the Cambridge Mathematics Placements (CMP) programme.

