## Supplementary Information for "Dynamic, adaptive sampling during nanopore sequencing using Bayesian experimental design"

April 1, 2022

**3 Supplementary figures** **23**

**4 Supplementary tables** **32**

**References** **34**

### 1 Supplementary methods

#### 1.1 Defining priors of genotype probabilities and updating posteriors after observing sequencing reads

**Priors of genotype probabilities** In the simplest case of a haploid genome without indels, we define the prior of reference nucleotide $b_R$ with $b_R \in B$ and $B = \{\mathrm{A}, \mathrm{C}, \mathrm{G}, \mathrm{T}\}$ at position $i$ as $\pi_i(b_R) = 1-\theta$, with $\theta$ the genetic diversity of the considered population. Conversely, $\pi_i(g) = \theta/3$ if $g \neq b_R$, with $g \in G$ and $G = B$ in this case.

When considering diploid sequenced genomes, we still assume a haploid reference genome, with reference nucleotide at a given position denoted $b_R$. And equivalently to the main text the set of possible genotypes $G$ instead consists of the unordered pairs $g = \{b_1, b_2\}$, with $b_1, b_2 \in B$. For an unphased genome without indels, we define $\pi_i(\{b_R, b_R\}) = 1-\theta$, and $\pi_i(\{g, g\}) = p_{\text{homo}}\theta/3$ if $g \neq b_R$, with $p_{\text{homo}}$ being the proportion of site differences from a reference that are expected to be homozygous, and $\pi_i(\{g, b_R\}) = (1 - p_{\text{homo}})\theta/3$ for $g \neq b_R$.

**Updating posteriors** In the main text we showed how to calculate the posterior probability $f_i(g|D)$ of genotypes $g$ at position $i$, conditional on data $D$ (Eq. 1). For completeness, here we provide details on the observation probabilities $\phi$, and further details on how to update the posterior distribution after observing additional data.

First, if observed data $D$ contains $n$ reads covering position $i$ with bases $d_{j,i}$ for $j = 1 \dots n$, we denote the base from a new hypothetical read at position $i$ by $d_{n+1,i}$. We then represent $D'$ as the union of $D$ with the new hypothetical read, so that $D'$ contains $n + 1$ reads covering position $i$, with bases $d_{j,i}$ for $j = 1 \dots n + 1$. After observing the new read we update the prior genotype probabilities $\pi_i(g)$ to get the posterior probabilities:

$$\begin{aligned} f_i(g|D') &= \frac{\pi_i(g) \prod_{j=1}^{n+1} \phi(d_{j,i}|g)}{\sum_{c \in G} \left( \pi_i(c) \prod_{j=1}^{n+1} \phi(d_{j,i}|c) \right)} \\ &= \frac{f_i(g|D) Z_i(D) \phi(d_{n+1,i}|g)}{\sum_{c \in G} f_i(c|D) Z_i(D) \phi(d_{n+1,i}|c)} \\ &= \frac{f_i(g|D) \phi(d_{n+1,i}|g)}{\sum_{c \in G} f_i(c|D) \phi(d_{n+1,i}|c)} \,. \end{aligned} \tag{S.1}$$

As in the main text, $Z_i(D)$ represents a normalising constant ensuring that the posterior probabilities at site $i$ sum to 1. $\phi(d_{j,i}|g)$ is the probability of calling base $d_{j,i}$ assuming genotype $g$ at position $i$, and will depend on assumptions about the probabilities of observing errors. For example, for a haploid genome without indels we define

$$\phi(d_{j,i}|b) = \begin{cases} 1 - e\,, & \text{if } d_{j,i} = b \in B\,, \\ \dfrac{e}{3}\,, & \text{if } d_{j,i} \neq b \in B\,, \end{cases} \tag{S.2}$$

where $e$ denotes the per-base substitution error probability, i.e. any position in a read has probability $e$ of representing a wrong nucleotide. In the scenario of an unphased diploid genome without indels we instead consider

$$\phi(d_{j,i}|\{b_1, b_2\}) = \begin{cases} 1 - e\,, & \text{if } d_{j,i} = b_1 = b_2\,, \\ \dfrac{1-e}{2} + \dfrac{e}{6}\,, & \text{if } d_{j,i} = b_1 \neq b_2 \text{ or } d_{j,i} = b_2 \neq b_1\,, \\ \dfrac{e}{3}\,, & \text{if } d_{j,i} \neq b_1,\ b_2\,. \end{cases} \tag{S.3}$$

#### 1.2 Incorporating deletions in the model

In this section we discuss how deletions, appearing either as mutational events or sequencing errors, are incorporated into our framework. Insertions and rearrangements are not considered due to increased complexity. If we include deletions the possible set of observed bases for haploid genomes becomes $B = \{\mathrm{A}, \mathrm{C}, \mathrm{G}, \mathrm{T}, -\}$, with the genotypes $G = B$. For an unphased diploid genome, $g \in G$ instead becomes one of 15 unordered pairs $g = \{b_1, b_2\}$, with $b_1, b_2 \in B$.

In order to define priors on these genotypes, we use parameter $r$ to express how often variation in the form of a deletion is observed relative to SNPs. Across a haploid sequenced genome we expect $N\theta$ substitutions from the reference and $rN\theta$ deleted bases. In our experiments we use values $\theta = 0.01$ and $r = 0.4$, in line with values reported for human populations and microbiomes (Schloissnig et al. 2013; The 1000 Genomes Project Consortium 2015).

Taking deletions into account requires modification of prior genotype probabilities $\pi$ and sequencing probabilities $\phi(d_{j,i}|g)$. For a haploid genome the prior of reference nucleotide $b_R$ at position $i$ is $\pi_i(b_R) = 1 - \theta(1 + r)$. Conversely, $\pi_i(g) = \theta/3$ if $g \neq b_R, -$; otherwise $\pi_i(-) = r\theta$.

For a diploid unphased genome, we define

$$\pi_i(\{g_1, g_2\}) = \begin{cases} 1 - \theta(1 + r)\,, & \text{if } g_1 = g_2 = b_R\,, \\ p_{\text{homo}}\theta/3\,, & \text{if } g_1 = g_2 \neq b_R, -\,, \\ (1 - p_{\text{homo}})\theta/3\,, & \text{if } g_1 = b_R \text{ and } g_2 \neq b_R, -\,, \\ (1 - p_{\text{homo}})\theta/3\,, & \text{if } g_2 = b_R \text{ and } g_1 \neq b_R, -\,, \\ p_{\text{homo}}r\theta\,, & \text{if } g_1 = g_2 = -\,, \\ (1 - p_{\text{homo}})r\theta\,, & \text{if } g_1 = - \text{ and } g_2 = b_R\,, \\ (1 - p_{\text{homo}})r\theta\,, & \text{if } g_2 = - \text{ and } g_1 = b_R\,. \end{cases} \tag{S.4}$$

Sequencing probabilities $\phi$ are modified in the following way. As before, $e$ represents the error probability for substitutions, and in all applications we use $e = 0.06$. We also define $e_-$ as the probability of a deletion sequencing error, i.e. observing a deletions instead of a base, set to $e_- = 0.05$ in our applications; finally, $e_+$ represents the probability that an actual deleted base in the sequenced genome is misread as a nucleotide, set to $e_+ = 0.1$ in our applications. For a haploid genome the sequencing probabilities then become

$$\phi(d_{j,i}|b) = \begin{cases} 1-(e+e_-)\,, & \text{if } d_{j,i}=b\,,\ b\neq -\,,\\ \dfrac{e}{3}\,, & \text{if } d_{j,i}\neq b\,,\ b\neq -\,,\ d_{j,i}\neq -\,,\\ e_-\,, & \text{if } d_{j,i}=-\,,\ b\neq -\,,\\ 1-e_+\,, & \text{if } d_{j,i}=b=-\,,\\ \dfrac{e_+}{4}\,, & \text{if } d_{j,i}\neq b\,,\ b=-\,. \end{cases} \tag{S.5}$$

In the scenario of an unphased diploid genome we instead have:

$$\phi(d_{j,i}|\{b_1,b_2\}) = \begin{cases} 1-(e+e_-)\,, & \text{if } d_{j,i}=b_1=b_2\neq -\,,\\ \dfrac{1-(e+e_-)}{2}+\dfrac{e}{6}\,, & \text{if } -\neq b_1,b_2,\ \text{ and } b_1\neq b_2,\ \text{ and } d_{j,i}\in\{b_1,b_2\}\,,\\ \dfrac{e}{3}\,, & \text{if } d_{j,i}\neq b_1\,,\ b_2\,,\ -\text{ and } b_1\,,\ b_2\neq -\,,\\ e_-\,, & \text{if } d_{j,i}=-\neq b_1\,,\ b_2\,,\\ 1-e_+\,, & \text{if } d_{j,i}=b_1=b_2=-\,,\\ \dfrac{e_+}{4}\,, & \text{if } d_{j,i}\neq b_1=b_2=-\,,\\ \dfrac{e_-}{2}+\dfrac{1-e_+}{2}\,, & \text{if } d_{j,i}=b_1=-\neq b_2\,,\\ \dfrac{1-(e+e_-)}{2}+\dfrac{e_+}{8}\,, & \text{if } b_1=-\neq b_2=d_{j,i}\,,\\ \dfrac{e}{6}+\dfrac{e_+}{8}\,, & \text{if } d_{j,i}\,,\ b_2\neq b_1=-\text{ and } d_{j,i}\neq b_2\,. \end{cases} \tag{S.6}$$

#### 1.3 Calculating KL-divergence to express positional site-wise score

In the main text we use the Kullback-Leibler divergence between the posterior probability distributions before and after observing a new read ($f_i(g|D)$ and $f_i(g|D')$, respectively) at site $i$ as a measure of potential information gain, where $D$ denotes already observed data and $D'$ represents augmentation of the data by one sequencing read, $d_{n+1,i}$. Additionally, we take the probability of observing any base in that read into account (main text, Eqs. 2 and 3). Here we present a derivation and practical form for calculating the KL divergence between the two aforementioned posterior probability distributions. We give some examples of positional site-wise scores $S_i$ resulting from different coverage patterns in Suppl. Fig. 1.

$$
\begin{aligned}
S_i &= \sum_{d_{n+1,i}\in B} P(d_{n+1,i}|D) D_{\mathrm{KL}}(f_i(g|D') \,||\, f_i(g|D)) \\
&= \sum_{d_{n+1,i}\in B} P(d_{n+1,i}|D) \left( \sum_{g\in G} f_i(g|D') \log \frac{f_i(g|D')}{f_i(g|D)} \right) \\
&= \sum_{d_{n+1,i}\in B} \sum_{g\in G} P(d_{n+1,i}|D) f_i(g|D') \log f_i(g|D') \\
&\qquad - \sum_{g\in G} \log f_i(g|D) \left( \sum_{d_{n+1,i}\in B} P(d_{n+1,i}|D) f_i(g|D') \right) \\
&= \sum_{d_{n+1,i}\in B} \sum_{g\in G} P(d_{n+1,i}|D) f_i(g|D') \log f_i(g|D') \\
&\qquad - \sum_{g\in G} \log f_i(g|D) \left( \sum_{d_{n+1,i}\in B} P(d_{n+1,i}|g, D) f_i(g|D) \right) \\
&= \sum_{d_{n+1,i}\in B} \sum_{g\in G} P(d_{n+1,i}|D) f_i(g|D') \log f_i(g|D') \\
&\qquad - \sum_{g\in G} f_i(g|D) \log f_i(g|D) \left( \sum_{d_{n+1,i}\in B} P(d_{n+1,i}|g, D) \right) \\
&= \sum_{d_{n+1,i}\in B} \sum_{g\in G} P(d_{n+1,i}|D) f_i(g|D') \log f_i(g|D') - \sum_{g\in G} f_i(g|D) \log f_i(g|D)\,. \qquad \text{(S.7)}
\end{aligned}
$$

##### 1.4 Practical calculation of the expected benefit of sequencing reads

In the main text we defined a positional benefit score $S_i$ for each position $i$ of a genome, and combined the scores of multiple positions and the distribution of previously observed read lengths into an expected benefit of reads $U_i$, assuming that each read maps to a series of contiguous bases in the reference genome (see main text, Eq. 5).

As part of Eq. 5, we define $S^l_{i,1}$ to be the sum of $l$ consecutive $S_j$ score values starting at position $i$, that is, the score of a forward-oriented read of length $l$ starting at position $i$:

$$S^l_{i,1} = \sum_{j=i}^{i+l-1} S_j \, . \tag{S.8}$$

Similarly, for a reverse-oriented read:

$$S^l_{i,0} = \sum_{j=i-l+1}^{i} S_j \, . \tag{S.9}$$

Since we do not know the length of a sequencing read in advance we account for the uncertainty in $l$. For this, we assume a single distribution of fragment lengths that applies to all fragments irrespective of genomic origin or orientation. We denote the fragment length distribution by $L(l)$ for lengths $l = 1 \ldots N$, with mean $\lambda = \sum_{l=1}^{N} L(l)l$.

In our real-time applications we use a truncated normal distribution with parameters $\lambda = 6,000$ and $\text{sd} = 4,000$ as a prior. Throughout the sequencing experiment this prior distribution is updated by the observed read lengths of full-sized, accepted reads to guarantee accurate calculation of the expected benefit of sequencing reads. This is especially important when targeting sparsely distributed variant sites, due to the potential information gained when a read might cover them. We observed that this adaptive, empirical read length distribution is learned within the first minutes of sequencing.

Since there will be lower and upper limits on the length of fragments, it is computationally convenient to define $\mathscr{D}_L$ to be the domain of $L$, i.e. the set of values of $l$ with $L(l) > 0$.

For more convenient calculation of equation 5 in the main text, we can use the cumulative distribution of read lengths, $CL(l) = \sum_{j=1}^{l} L(j)$, instead. Considering its corresponding complementary distribution $\widetilde{CL}\,(l) = 1 - CL(l) = \sum_{j=l+1}^{N} L(j)$, Eq. 5 can be rewritten as

$$U_{i,1} = \sum_{l \in \mathfrak{D}_{\widetilde{CL}}} \widetilde{CL}\,(l) S_{i+l-1}\,. \tag{S.10}$$

Analogously, for reverse reads:

$$U_{i,0} = \sum_{l \in \mathfrak{D}_{\widetilde{CL}}} \widetilde{CL}\,(l) S_{i+1-l}\,. \tag{S.11}$$

Calculating $U_{i,1}$ and $U_{i,0}$ for all genome positions with a naive algorithm would require time proportional to $|\mathfrak{D}_{\widetilde{CL}}|$. As $U_{i,1}$ needs to be calculated for each $i$, the total cost for the whole genome would be in the order of $O(N \times |\mathfrak{D}_{\widetilde{CL}}|)$, which would be excessively slow. We therefore efficiently and accurately approximate $U_i$, with total computational cost linear in genome size, using an approach based on approximating $\widetilde{CL}$ with a piece-wise constant function.

Assuming that $\widetilde{CL}\,(l)$ is a piece-wise constant function means that there are $\eta$ values $1 = x_1 < x_2 < \ldots < x_\eta = \max \mathfrak{D}_{\widetilde{CL}} + 1$ such that for all $1 \leq \nu < \eta$ and for all $x \in [x_\nu, x_{\nu+1})$ we have $\widetilde{CL}\,(x) = \widetilde{CL}\,(x_\nu)$. Equivalently to the case without approximation we calculate the benefit of the first position:

$$U_{1,1} = \sum_{l \in \mathfrak{D}_{\widetilde{CL}}} \widetilde{CL}\,(l) S_l\,. \tag{S.12}$$

This still requires the same time as discussed previously. However, calculating $U_{i,1}$ for every other genome position $i > 1$ now requires only time $O(\eta)$, where $\eta$ is the number of different constant values in $\widetilde{CL}$. In general, if we know $U_{i,1}$ we can calculate $U_{i+1,1}$ as

$$U_{i+1,1} = U_{i,1} - S_i + S_{i+x_\eta-1}\,\widetilde{CL}\,(x_{\eta-1}) + \sum_{2 \leq \nu < \eta} \left(\widetilde{CL}\,(x_{\nu-1}) - \widetilde{CL}\,(x_\nu)\right) S_{i+x_\nu-1}\,. \tag{S.13}$$

The same approach can be used for efficiently calculating the expected benefit of reverse reads ($U_{i,0}$). In our experiments we use a piece-wise constant function with $\eta = 11$ different values.

#### 1.5 Details on deriving the decision framework

In the main text we defined a strategy $\mathcal{S}$ to be a function $I^{\mathcal{S}}_{i,o}$ returning 0 or 1 for reads from any position $i$ in a genome with orientation $o$. Thus, $I^{\mathcal{S}}_{i,1} = 0$ indicates that a forward fragment starting at position $i$ should be rejected, while $I^{\mathcal{S}}_{i,0} = 1$ indicates that a reverse fragment starting at position $i$ should be read to its end, and so on. Here, we present some more details on the calculation of the decision strategy and some generalisations. We say that $\mathcal{S}$ *includes* $(i, o)$ if $I^{\mathcal{S}}_{i,o} = 1$. Since we do not know $\mathcal{S}$ *a priori*, our aim is to determine an optimal strategy $\widehat{\mathcal{S}}$ given the current data $D$.

Given the definitions in the main text and above, the expected benefit of a DNA fragment of orientation $o$ starting at position $i$ is

$$U^{\mathcal{S}}_{i,o} = S^{\mu}_{i,o} + I^{\mathcal{S}}_{i,o}\big(U_{i,o} - S^{\mu}_{i,o}\big) \,, \tag{S.14}$$

with $S^{\mu}_{i,o}$ denoting the expected benefit of the initial $\mu$ bases of a read and calculated according to Suppl. Eqs. S.8 and S.9. This accumulation of benefit is achieved in time

$$t^{\mathcal{S}}_{i,o} = \mu + I^{\mathcal{S}}_{i,o}(\lambda - \mu) + \big(1 - I^{\mathcal{S}}_{i,o}\big)\rho + \alpha = \alpha + \mu + \rho + I^{\mathcal{S}}_{i,o}\left(\lambda - \mu - \rho\right) \,. \tag{S.15}$$

**Accounting for bias in the origin of sequencing reads** In the main text we present simplified equations assuming a uniform probability for the origin of DNA fragments. In reality, however, calculating the strategy-wise average time cost $\bar{t}^{\mathcal{S}}$ and benefit $\bar{U}^{\mathcal{S}}$ requires knowledge about how often fragments from certain positions $i$ and orientation $o$ are sequenced by pores.

In many cases variation in this distribution could be ignored without much impact on the computed strategies, especially if there is little coverage bias, e.g. when sequencing input DNA from a single species without prior amplification. Sometimes however, ignoring these probabilities could negatively influence the optimality of the decision strategy. For example regions with very low coverage, i.e. negative coverage bias, will have high expected benefit but low probability of being covered by future reads. This could lead to over-rejection of fragments due to expecting lots of benefit from areas where reads are unlikely to originate in the future.

Accordingly, in our implementation we generalise and account for bias is the origin of sequencing reads. For this, we use the notation $F_{i,o}$ to refer to the probability of a random fragment's first base mapping to position $i$ in orientation $o$, so that $\sum_{o=1,0}\sum_{i=1}^{N} F_{i,o} = 1$. In this case, the average benefit per fragment $\bar{U}^{\mathcal{S}}$ of strategy $\mathcal{S}$ becomes

$$\begin{aligned}\bar{U}^{\mathcal{S}} &= \sum_{o=1,0}\sum_{i=1}^{N} F_{i,o} U_{i,o}^{\mathcal{S}} \\ &= \sum_{o=1,0}\sum_{i=1}^{N} F_{i,o}\left(S_{i,o}^{\mu} + I_{i,o}^{\mathcal{S}}\big(U_{i,o} - S_{i,o}^{\mu}\big)\right) \end{aligned} \tag{S.16}$$

and its average fragment-wise cost $\bar{t}^{\mathcal{S}}$ is given by

$$\begin{aligned}\bar{t}^{\mathcal{S}} &= \sum_{o=1,0}\sum_{i=1}^{N} F_{i,o} t_{i,o}^{\mathcal{S}} \\ &= \alpha + \mu + \rho + (\lambda - \mu - \rho)\sum_{o=1,0}\sum_{i=1}^{N} F_{i,o} I_{i,o}^{\mathcal{S}} \,. \end{aligned} \tag{S.17}$$

Note that in the simplest case, with no bias in read origin or orientation, $F_{i,o} = 1/2N$ for each of the $2N$ position-orientation pairs $(i,o)$. In this case, we can write $\bar{S}^{\mu} = (\sum_{o=1,0}\sum_{i=1}^{N} S_{i,o}^{\mu})/2N$.

Modeling and updating of the distribution of read origins $F_{i,o}$ is further detailed in Suppl. Sect. 1.6.

**Formalised procedure to find optimal strategy** To find optimal strategies, we first rank all the $2N$ position-orientation pairs $(i,o)$ according to decreasing value of $U_{i,o} - S_{i,o}^{\mu}$ and index them such that $(i_1, o_1)$ takes the highest value, $(i_2, o_2)$ the next and so on: $U_{i_1,o_1} - S_{i_1,o_1}^{\mu} \geq U_{i_2,o_2} - S_{i_2,o_2}^{\mu} \geq \ldots \geq U_{i_{2N},o_{2N}} - S_{i_{2N},o_{2N}}^{\mu}$.

Strategy $\mathcal{S}^{\sigma}$ is defined by setting $I_{\imath}^{\mathcal{S}^{\sigma}} = 1$ for $\imath = (i_1, o_1)\ldots(i_{\sigma}, o_{\sigma})$ and 0 otherwise, and it is the optimal strategy of size $\sigma$. Starting with $\sigma = 0$, the empty strategy, we successively increase $\sigma$, at each stage testing whether

$$\frac{U_{i_{\sigma+1},o_{\sigma+1}} - S_{i_{\sigma+1},o_{\sigma+1}}^{\mu}}{\lambda - \mu - \rho} > \frac{\bar{U}^{\mathcal{S}^{\sigma}}}{\bar{t}^{\mathcal{S}^{\sigma}}} \tag{S.18}$$

to discover whether $\mathcal{S}^{\sigma+1}$ gives an improvement over $\mathcal{S}^{\sigma}$. Once we reach a value $\sigma^*$ such that there is no further improvement, we have the optimal $\widehat{\mathcal{S}} = \mathcal{S}^{\sigma^*}$.

##### 1.6 Accounting for variation in the distribution of sequencing reads

In Suppl. Sect. 1.5 we show how we account for variation in the origin of sequenced fragments by incorporating a distribution $F_{i,o}$. In most scenarios simply incorporating observed read starting positions in an empirical distribution would suffice. However, there are other occasions when this is not enough: specifically, when parts of the reference genome are not present in the sequenced sample, e.g. when sequencing diverged bacterial strains, some regions might not receive any coverage at all and would continuously dominate the ranking of yet-to-be-gained expected benefit.

We thus model variation in $F_{i,o}$ and estimate it from currently observed data using a Bayesian approach. Since the $F_{i,o}$ define a discrete multinomial probability distribution, it makes sense to choose a Dirichlet distribution, its conjugate prior, as the prior over the $F_{i,o}$ parameters. We expect, however, a non-negligible proportion of $F_{i,o}$ to be exactly zero (sites at which no read's mapping starts, for example due to deletions relative to the reference). Therefore, we use a mixed distribution of a point mass at 0 and a Dirichlet distribution as a prior for $F_{i,o}$.

In the approach discussed below the positional index $i$ can refer equally to individual genomic positions or windows of consecutive sites. That way, $F_{i,o}$ refers to windows rather than individual positions, reducing the variance of the estimate and required computational resources.

We assume parameter $a_{i,o} = a$ for each position-orientation pair $(i, o)$ in the Dirichlet prior and represent the distribution as $\text{Dir}(a_{1,0}, \ldots, a_{N,0}, a_{1,1}, \ldots, a_{N,1}) = \text{Dir}(a)$. To define the value of $a$ used in practice, we use information from previous sequencing runs, e.g. observed variation in the empirical distribution of mapping positions. We define $C_{i,o}$ to be the count of reads starting at position $i$ and with orientation $o$, continuously updating these counts throughout a sequencing run. We can then define $\widehat{F}_{i,o} = C_{i,o} / \sum_{j,u} C_{j,u}$ as an approximate inference of the $F_{i,o}$, with variance $\widehat{V}$. An approximate estimate for $a$ is then found by equating $\widehat{V}$ to the variance of $\text{Dir}(a)$, i.e. $(2N-1)/4N^2(2Na+1)$. Rearranging the terms, this becomes

$$a = \frac{2N-1}{8N^3\widehat{V}} - \frac{1}{2N}. \tag{S.19}$$

Since we assume a mixture of point mass at 0 and a Dirichlet distribution, we have that any $F_{i,o}$ has a marginal prior distribution $P(F_{i,o} = p)$ given by

$$P(F_{i,o} = p) = \begin{cases} p_0\,, & \text{if } p = 0\,, \\ (1-p_0)\beta(a,(2N-1)a)\partial p\,, & \text{if } p > 0\,, \end{cases} \tag{S.20}$$

where $\partial p$ is an abuse of notation representing a differential in $p$: that is, while the probability of $F_{i,o} = 0$ is equal to $p_0$, the probability of $F_{i,o} = p$ for any $0 < p \leq 1$ is 0 because on $p > 0$ the prior is continuously distributed with a beta distribution (the marginal distribution of the Dirichlet distribution) with parameters $a$ and $(2N-1)a$, and rescaled by $1-p_0$. In other words, conditional on $F_{i,o} > 0$ we have $F_{i,o} \sim \beta(a,(2N-1)a)$.

Analogously to $a$, we use previously observed information to define $p_0$ and approximate it by the number of pairs $(i,o)$ that have counts $C_{i,o} = 0$ (i.e. no reads observed with that starting location and orientation).

Now that we have defined priors, we can calculate estimates $\widehat{F}_{i,o}$ of each $F_{i,o}$, using counts $C_{i,o}$. For simplicity and computational tractability, we estimate one $F_{i,o}$ at a time rather than trying to estimate all $F_{i,o}$ simultaneously; this is akin to using a composite likelihood approximation (Varin et al. 2011). We assume a binomial likelihood for $C_{i,o}$ given $F_{i,o}$, and use the posterior expected value of $F_{i,o}$ given $C_{i,o}$ as the estimate $\widehat{F}_{i,o}$. In the case of $C_{i,o} > 0$ the posterior expected value of $F_{i,o}$ is:

$$\begin{aligned} E[F_{i,o}|C_{i,o} > 0] &= \frac{(1-p_0)\int_{F_{i,o}>0} \beta(F_{i,o}|a,(2N-1)a)P(C_{i,o}|F_{i,o})F_{i,o}\partial F_{i,o}}{(1-p_0)P(C_{i,o}|F_{i,o} > 0)} \\ &= \frac{\int_{F_{i,o}>0} \beta(F_{i,o}|a,(2N-1)a)P(C_{i,o}|F_{i,o})F_{i,o}\partial F_{i,o}}{P(C_{i,o}|F_{i,o} > 0)} \\ &= \frac{a + C_{i,o}}{2Na + \sum_{j,u} C_{j,u}}\,, \end{aligned} \tag{S.21}$$

where $\beta(F_{i,o}|a,(2N-1)a)$ is the prior beta density function for $F_{i,o} > 0$ and $P(C_{i,o}|F_{i,o} > 0)$ is the probability of observing counts $C_{i,o}$ if we assume that $F_{i,o} > 0$, and therefore that $F_{i,o}$ has a simple beta density.

Updating the posterior expectation is more complicated with sites where $C_{i,o} = 0$. In this case $\widehat{F}_{i,o}$ is:

$$E[F_{i,o}|C_{i,o}=0] = \frac{(1-p_0)\int_{F_{i,o}>0}\beta(F_{i,o}|a,(2N-1)a)P(C_{i,o}=0|F_{i,o})F_{i,o}\partial F_{i,o}}{P(C_{i,o}=0)}$$
$$= \frac{(1-p_0)\int_{F_{i,o}>0}\beta(F_{i,o}|a,(2N-1)a)P(C_{i,o}|F_{i,o})F_{i,o}\partial F_{i,o}}{p_0+(1-p_0)P(C_{i,o}|F_{i,o}>0)} \tag{S.22}$$

As before, we substitute $\int_{F_{i,o}>0}\beta(F_{i,o}|a,(2N-1)a)P(C|F_{i,o})F_{i,o}\partial F_{i,o}$ with $P(C_{i,o}|F_{i,o}>0)(a+C_{i,o})/(2Na+\sum_{j,u}C_{j,u})$, due to the fact that $P(C|F_{i,o})$ is a binomial likelihood and that we are restricting ourselves to a beta density for $F_{i,o}$. The above equation then becomes:

$$E[F_{i,o}|C_{i,o}=0] = \frac{(1-p_0)P(C_{i,o}=0|F_{i,o}>0)}{p_0+(1-p_0)P(C_{i,o}=0|F_{i,o}>0)}\frac{a}{2Na+\sum_{j,u}C_{j,u}}$$
$$= \left(1-\frac{p_0}{p_0+(1-p_0)P(C_{i,o}=0|F_{i,o}>0)}\right)\frac{a}{2Na+\sum_{j,u}C_{j,u}}. \tag{S.23}$$

The only term above that needs further derivation is $P(C_{i,o}=0|F_{i,o}>0)$. Using the definition of density functions of the beta and binomial distributions, this is equal to:

$$P(C_{i,o}=0|F_{i,o}>0) = \frac{\int_{F_{i,o}>0}F_{i,o}^{a-1}(1-F_{i,o})^{(2N-1)a-1}(1-F_{i,o})^{\sum_{j,u}C_{j,u}}}{\mathrm{B}(a,(2N-1)a)}$$
$$= \frac{\mathrm{B}(a,(2N-1)a+\sum_{j,u}C_{j,u})}{\mathrm{B}(a,(2N-1)a)}, \tag{S.24}$$

where $\mathrm{B}(a,b)$ is the typical normalizing factor of the beta distribution density $\beta(a,b)$, that is, $\mathrm{B}(a,b)=\Gamma(a)\Gamma(b)/\Gamma(a+b)$, and where $\Gamma$ is the Gamma function $\Gamma(a)=\int x^{a-1}e^{-x}dx$. To calculate $\mathrm{B}(a,b)$ we do not use the Gamma function directly due to potential overflow, but instead use the natural logarithm of the beta function to get the logarithm of B. Finally, the estimators $\widehat{F_{i,o}}$ are normalised to ensure summation to 1.

In some scenarios, for example when sequencing only certain species from within a metagenomic sample, some reads will not map to the reference genomes considered. In such cases, we additionally account for the frequency of fragments in the input material that are not of interest. Otherwise, if a species of interest constitutes a low percentage of the input DNA, our model would overestimate the expected benefit of a new read and reject more reads than it should ideally. To circumvent this issue, we estimate the frequency of reads not mapping onto the reference by recording the number of on- and off-target reads and use that ratio for normalisation of the the estimators $\widehat{F}_{i,o}$.

##### 1.7 Proof of strategy optimality

Here we prove the results stated in the main text regarding the optimality of the fragment selection strategies proposed. We start by presenting two very general algebraic results that come in handy, represented here in terms of the variables we will use later in our proof. Assuming $u_\jmath,\ t_\jmath,\ U,\ T,\ u_\imath,\ t_\imath > 0$, we have that

$$
\begin{aligned}
\frac{u_\jmath}{t_\jmath} > \frac{U}{T}\ ,\ \frac{u_\imath}{t_\imath} &\iff u_\jmath T > t_\jmath U\ ,\ u_\jmath t_\imath > t_\jmath u_\imath \\
&\Rightarrow u_\jmath T + u_\jmath t_\imath > t_\jmath U + t_\jmath u_\imath \\
&\iff u_\jmath (T + t_\imath) > t_\jmath (U + u_\imath) \\
&\iff \frac{u_\jmath}{t_\jmath} > \frac{U + u_\imath}{T + t_\imath}
\end{aligned}
\tag{S.25}
$$

and

$$
\begin{aligned}
\frac{U + u_\imath}{T + t_\imath} \geq \frac{U}{T} &\iff T(U + u_\imath) \geq U(T + t_\imath) \\
&\iff T u_\imath \geq U t_\imath \\
&\iff \frac{u_\imath}{t_\imath} \geq \frac{U}{T}\ .
\end{aligned}
\tag{S.26}
$$

Now returning to optimal strategies, for greater generality we consider the case of multiple distinct reference chromosomes, potentially from different species. The only change needed is to refer to locations within a reference as $k, i$ to indicate their chromosome identifier $k$ as well as nucleotide location $i$; consequently, DNA fragments that are candidates for sequencing are referred to by triplets $(k, i, o)$ that indicate their chromosome and starting position within it, and orientation $o$. Now, we assume that positions $(k, i, o)$ have been ranked in an ordered list $\imath_1 \ldots \imath_{2N}$ such that $\imath_1$ has the highest value of $u_{k,i,o}/t_{k,i,o} = (U_{k,i,o} - S^{\mu}_{k,i,o})/(\lambda_{k,i,o} - \mu_{k,i,o} - \rho)$, $\imath_2$ has the second highest value, and so on. The only exception to this are the positions $(k, i, o)$ for which $t_{k,i,o} \leq 0$, which we assume have been ranked first or, equivalently, their score set to $u_{k,i,o}/t_{k,i,o} = +\infty$.

We define a strategy $\boldsymbol{\mathcal{S}}$ to be better than strategy $\boldsymbol{\mathcal{S}}'$ (represented as $\boldsymbol{\mathcal{S}} \succ \boldsymbol{\mathcal{S}}'$) if and only if it has greater expected benefit per unit time: $\bar{U}^{\boldsymbol{\mathcal{S}}}/\bar{t}^{\boldsymbol{\mathcal{S}}} > \bar{U}^{\boldsymbol{\mathcal{S}}'}/\bar{t}^{\boldsymbol{\mathcal{S}}'}$. Next, we show that the best strategy is one that accepts positions $\imath_s$ with $s \leq s^*$ and rejects positions $\imath_s$ with $s > s^*$ for some value of $s^*$. We ignore for simplicity the effects of positions with equal rank and scores. We use *reductio ad absurdum*, starting from the assumption that the best strategy includes a position $\imath$ but not a position $\jmath$ with $u_\jmath/t_\jmath > u_\imath/t_\imath$. If we denote the expected per-fragment benefit of this strategy but excluding $\imath$ by $U$, and the corresponding expected time by $T$, then the expected value of the best strategy is $(U+u_\imath)/(T+t_\imath)$. Then $(U+u_\imath)/(T+t_\imath) \geq U/T$ by the assumption of optimality and so $u_\imath/t_\imath \geq U/T$ (Eq. S.26); $u_\jmath/t_\jmath > u_\imath/t_\imath$ by assumption and hence $u_\jmath/t_\jmath > U/T$; and thus $u_\jmath/t_\jmath > (U + u_\imath)/(T + t_\imath)$ (Eq. S.25). This means that including position $\jmath$ would improve the best strategy (Eq. S.26 in reverse direction), which is a contradiction.

We now show that if we start from the null strategy $\boldsymbol{\mathcal{S}}^0$ (i.e. the strategy that rejects all fragments), successively add positions $\imath_1, \imath_2 \ldots$ (creating the series of strategies $\boldsymbol{\mathcal{S}}^1$, $\boldsymbol{\mathcal{S}}^2 \ldots$), and stop at the first $s^*$ such that $\boldsymbol{\mathcal{S}}^{s^*} \succ \boldsymbol{\mathcal{S}}^{s^*+1}$, then $\boldsymbol{\mathcal{S}}^{s^*}$ is the optimal strategy. From above, we already know that one of the $\boldsymbol{\mathcal{S}}^s$ must be the best strategy, and obviously for each $s < s^*$ we have $\boldsymbol{\mathcal{S}}^{s^*} \succ \boldsymbol{\mathcal{S}}^s$. We only have to show that for each $\delta \geq 1$, $\boldsymbol{\mathcal{S}}^{s^*} \succ \boldsymbol{\mathcal{S}}^{s^*+\delta}$. This is true for $\delta = 1$ (definition of $s^*$), so thanks to Eq. S.26 we have $U^{s^*}/T^{s^*} > u_{\imath_{s^*+1}}/t_{\imath_{s^*+1}}$, where we represent the expected benefit and cost per fragment of strategy $\boldsymbol{\mathcal{S}}^{s^*}$ as $U^{s^*} = \bar{S}^\mu + \sum_{s=1}^{s^*} u_{\imath_s}$ and $T^{s^*} = \alpha + \mu + \rho + \sum_{s=1}^{s^*} t_{\imath_s}$, respectively. By definition we have $U^{s^*+\delta} = U^{s^*} + \sum_{j=1}^{\delta} u_{\imath_{s^*+j}}$ and $T^{s^*+\delta} = T^{s^*} + \sum_{j=1}^{\delta} t_{\imath_{s^*+j}}$, and therefore

$$
\begin{aligned}
\boldsymbol{\mathcal{S}}^{s^*} \succ \boldsymbol{\mathcal{S}}^{s^*+\delta} &\iff \frac{U^{s^*}}{T^{s^*}} > \frac{U^{s^*} + \sum_{j=1}^{\delta} u_{\imath_{s^*+j}}}{T^{s^*} + \sum_{j=1}^{\delta} t_{\imath_{s^*+j}}} \\
&\iff \frac{U^{s^*}}{T^{s^*}} > \frac{\sum_{j=1}^{\delta} u_{\imath_{s^*+j}}}{\sum_{j=1}^{\delta} t_{\imath_{s^*+j}}} \qquad \text{(S.27)}
\end{aligned}
$$

with the last step coming from Eq. S.26. Since we know that $U^{s^*}/T^{s^*} > u_{\imath_{s^*+1}}/t_{\imath_{s^*+1}}$, it is sufficient to prove that for any $\delta \geq 1$ we have $u_{\imath_{s^*+1}}/t_{\imath_{s^*+1}} \geq \sum_{j=1}^{\delta} u_{\imath_{s^*+j}} / \sum_{j=1}^{\delta} t_{\imath_{s^*+j}}$. This is obviously true for $\delta = 1$, and we use the induction principle to prove it for any $\delta \geq 2$. Inductively, assuming that $u_{\imath_{s^*+1}}/t_{\imath_{s^*+1}} \geq (\sum_{j=1}^{\delta-1} u_{\imath_{s^*+j}})/(\sum_{j=1}^{\delta-1} t_{\imath_{s^*+j}})$ then from the fact that $u_{\imath_{s^*+1}}/t_{\imath_{s^*+1}} \geq u_{\imath_{s^*+\delta}}/t_{\imath_{s^*+\delta}}$ and from Eq. S.25 we obtain the required thesis.

#### 1.8 Efficiency improvements required to ensure strategy optimality

To ensure optimality of dynamic sequencing strategies we require an efficient algorithm that can update the strategy in short intervals in order to keep up with the data stream from the sequencing device. One of the bottlenecks is ranking sites by their expected benefit. We therefore conceived a fast algorithm based on approximating the expected benefit of reads by discretized values. To do this, we decompose the floating point benefit values into their significand and exponent components, and use the exponents to form a grid approximation while ignoring the significands. This way, instead of sorting $2N$ floats (for total reference genome size $N$), we tally the counts of integers, which is easily parallelizable. Consequently, this also means that for finding the size $\sigma$ of the decision strategy, i.e. the number of ranked positions from which to accept reads, we do not operate on a per-site basis but instead test for an improvement of our optimality criterion after adding the multiple sites from one point of the discretised grid. Therefore, we reduce the number of considered instances from $2N$ sites to the number of points in the grid approximation. In reality the truly optimal threshold will likely lie between two points on the grid, so as a trade-off for computational speed we are likely either accepting or rejecting reads from few additional sites compared to a strategy calculated with an exact approach. This algorithm is described in Algorithm 1.

This procedure speeds up the calculation of new strategies but is still computationally impractical when considering large genomes or multiple species in a single experiment. In such settings, the real-time data stream from the sequencing device could outpace the generation of new decision strategies. To prevent this, we use a further approximation based on the assumption that neighboring sites will often have very similar values of expected benefit. This is justified by the fact that reads starting at some position will have a high probability of covering very similar consecutive sites as a read with the same orientation starting at a neighbouring site. Therefore, we reduce the resolution of the generated decision strategy by taking the sum of positional benefit scores in non-overlapping windows of size $w$, i.e. we calculate exact posterior probabilities and scores for each site, but obtain values of expected benefit of reads starting within a window of $w$ bases instead of calculating the benefit of reads starting at any position $i$. Subsequently, we also generate decisions for windows of $w$ bases instead of decisions for individual positions, i.e. we might use index $w$ in the indicator function $I^{\mathcal{S}}_{w,o}$ to describe whether a read from that window

**Algorithm 1: Finding approximate decision strategies.** By using a grid approximation, we consider whether reads from points on the grid, i.e. all sites at one point of the grid instead of individual sites, should be added to the strategy. Since the number of grid points considered is typically much smaller than the number of sites, this leads to a significant reduction in the number of comparisons needed, in addition to avoiding the step of sorting a vector of $2N$ values.

**Input:** vector $U$ of expected benefits per site; vector $t$ of sequencing time cost per site
**Output:** decision strategy $\widehat{\mathcal{S}}$

$U' \leftarrow$ scale expected benefit $U$ by $\max(U)$
$S, E \leftarrow$ discretize benefit by decomposing $U'$ into significands and exponents
$G \leftarrow$ dictionary storing counts of points in the grid approximation, with keys $k$
**for** $e$ *in* $E$ **do** `// count occurrences to form grid approx.`
  $G_{|e|} \leftarrow G_{|e|} + 1$
**end**
$U^G \leftarrow$ vector for expected benefit of grid approximation
$K \leftarrow$ sorted vector of grid points (keys $k$ of $G$)
**for** $k$ *in* $K$ **do** `// recreate approx. expected benefit of points in grid`
  $U^G_k \leftarrow 2^k \ \max(U)$
**end**
$\bar{U}^0 \leftarrow$ average benefit if all fragments are rejected (mean benefit of initial, length $\mu$ fragment)
$\bar{t}^0 \leftarrow$ average time cost if all fragments are rejected (time to acquire fragment, sequence $\mu$ bases and make decision)
$t' \leftarrow$ average time cost of sites at grid point
**for** $k$ *in* $K$ **do**
  $\bar{U}^k \leftarrow \sum_{i=1}^{k} U^G_i \ G_i + \bar{U}^0$ `// cumulative sum of benefit and grid counts`
  $\bar{t}^k \leftarrow \sum_{i=1}^{k} t'_i \ G_i + \bar{t}^0$ `// cumulative sum of time cost and grid counts`
**end**
$\Theta \leftarrow \arg\max(\bar{U}/\bar{t})$ `// threshold to maximise rate of benefit per time unit`
$\widehat{\mathcal{S}} \leftarrow$ find where $U \geq \Theta$

should be accepted or rejected. In our applications we select $w = 100$ to achieve a speed-up of two orders of magnitude, while keeping the resolution of the decisions well below commonly observed read lengths in order not to negatively affect optimality. If reads in an input library are expected to be especially long, $w$ could possibly be set to larger values to achieve even greater performance gains.

Another way of decreasing computational cost at runtime is to precompute the positional benefit scores $S_i$ of the most commonly observed coverage patterns that we expect to observe during a sequencing run; that is, patterns of counts of observed bases $b \in B$ amongst the $n$ reads, $d_{j,i}$ for $j = 1 \ldots n$, at position $i$. For this, we generate various combinations of counts of the reference and non-reference alleles as well as deletions and calculate the posterior probabilities for genotypes and resulting positional scores of the coverage patterns. By default we consider up to 40× coverage as no significant change in the genotype probability distribution is expected after that amount of data has been observed at a site. The precomputed values are stored in a multidimensional array that is indexed by the coverage counts of nucleotides and deletions, enabling very fast retrieval. Precomputing the most commonly encountered coverage patterns means that we only need to compute posterior probabilities and positional benefit scores for sites with unusual coverage counts during runtime, e.g. sites with unexpected allelic proportions or increased sequencing error.

### 2 Supplementary results

#### 2.1 Sequencing ROIs of two species in the presence of abundance bias

In the main text we present a sequencing experiment of a microbial mixture where we are interested in the entire genome of every species. In contrast, we might only want to investigate a smaller proportion of a few genomes in a community. This could be, for example, in order to quickly interrogate the presence of antimicrobial resistance-associated (AMR) loci in a clinical setting. We chose to use the same microbial mock community with logarithmically distributed abundances (ZymoBIOMICS DNA Standard II D6311, Zymo Research), but focused our sequencing effort on AMR loci of the two most abundant species, *L. monocytogenes* and *P. aeruginosa*. For this, we employed the same closely related but not identical reference genomes as described in the main text and used them to inform priors for the genotype probability distribution. Before sequencing we identified AMR loci using the CARD database (Alcock et al. 2020), resulting in ROIs covering 12.4% and 9.8% of the bacterial genomes, respectively. We added 10kb of flanking regions ahead of each ROI to account for reads that start close enough to cover them. This resulted in initial strategy sizes of 64.7% and 57.9% of the genome for *P. aeruginosa* and *L. monocytogenes*, respectively.

In this experiment we did not only compare BOSS-RUNS to sequencing without using adaptive sampling, but also to readfish. Readfish is an established tool that is well-suited to target ROIs in genomes. With this comparison we aim to highlight the additional benefit of dynamically adjusted adaptive sampling in acquiring more sequencing data at the most relevant positions, e.g. at regions within genomes affected by coverage bias.

As with the application presented in the main text, sequencing was conducted on a GridION using R9.4 flowcells. Out of 512 total channels on the flowcell each of the three conditions was assigned 128 sequencing channels. Readfish was configured to reject reads if they were found to map to one or more off-target sites, i.e. to sites outside of the specified ROIs, or if they did not map at all or failed to produce reliable basecalls. Again, we used a data chunk of 0.8s as the initial fragment for inferring the genomic origin and orientation in order to make decisions.

As expected, the proportion of sites from which reads are accepted decreases very quickly for *L. monocytogenes*, followed by a more steady decline for the less abundant *P. aeruginosa* (Suppl. Fig. 7A). Interestingly, while the strategy size of *L. monocytogenes* is very small, there are still a few different sites at which the expected benefit is large enough to warrant sequencing reads in their entirety despite the coverage difference of an order of magnitude. These sites include positions of substantial differences to the reference we used or sites which are inherently difficult to resolve, such as homopolymers (Suppl. Fig. 7A, inset).

The mean coverage depth of the two species demonstrates that we successfully trade-off additional data from *L. monocytogenes* in order to boost the coverage of *P. aeruginosa*. Especially in the first few hours of sequencing we achieve the highest coverage values for the rarer of the two species compared to both the control and readfish (Suppl. Fig. 7B).

The advantage of focusing data collection on ROIs with BOSS-RUNS compared to readfish becomes more evident, however, when we consider how the sequencing data is distributed within ROIs. Here, we observe that the proportion of sites that remain covered by less than 5× is lowest for BOSS-RUNS. Equally, the remaining total uncertainty, i.e. the remaining entropy of the genotype probability distributions across all sites of interest, is also lowest when using our new method (Suppl. Fig. 7C,D). In fact, BOSS-RUNS surpasses the total acquired information content of the control and readfish in the ROIs of *P. aeruginosa* after 27.1% and 43.0% of the sequencing run, respectively.

Visualising the entire distribution of coverage depth at multiple time points throughout the experiment is another way to show how BOSS-RUNS redistributes data within species. In Suppl. Fig 8 we compare the collected coverage of the control sector (A) to BOSS-RUNS (B) and readfish (C). Readfish achieves the largest enrichment of data as shown by the mean coverage depth of both species (Suppl. Fig. 7B) and the entire distribution (Suppl. Fig. 8C). BOSS-RUNS on the other hand does not continue to collect data from all of the specified ROIs, but instead stops sampling data from regions after observing enough data to resolve the genotypes and subsequently manages to enrich coverage at specific areas where it is needed most. This results not only in fewer sites at coverage <5×, but overall at distributions with lower variance (Suppl. Fig. 7B).

Analysing the enrichment of on-target versus off-target regions confirms previous observations. By normalising the total yield of sequencing to the control sector of the flowcell, we see that readfish collected 1.5× more data of on-target sites in both bacteria compared to the control and 0.6× the amount of data from the remaining parts of the genome (Suppl. Fig. 9). BOSS-RUNS instead only accumulates 50% of the amount of data in *L. monocytogenes* compared to the control as this bacteria's chromosome is quickly resolved. In the case of *P. aeruginosa*, BOSS-RUNS collects only slightly more data from ROIs but depletes off-target regions equally well as readfish. This smaller enrichment compared to readfish when measuring total yield is the trade-off for redistribution of data described previously. For this experiment we used the same input material and preparation methods as described in the main text, therefore resulting in a similar distribution of read lengths (see Fig. 2F). Since the amount of enrichment (and depletion) we can achieve is largely dependent on input read lengths, i.e. the expected difference between the initial fragment used to make decisions and the length of entire reads, the differences in yield presented here are well-suited to demonstrate the advantage of a dynamic aspect to adaptive sampling, but they could potentially be much larger given longer reads.

### 3 Supplementary figures

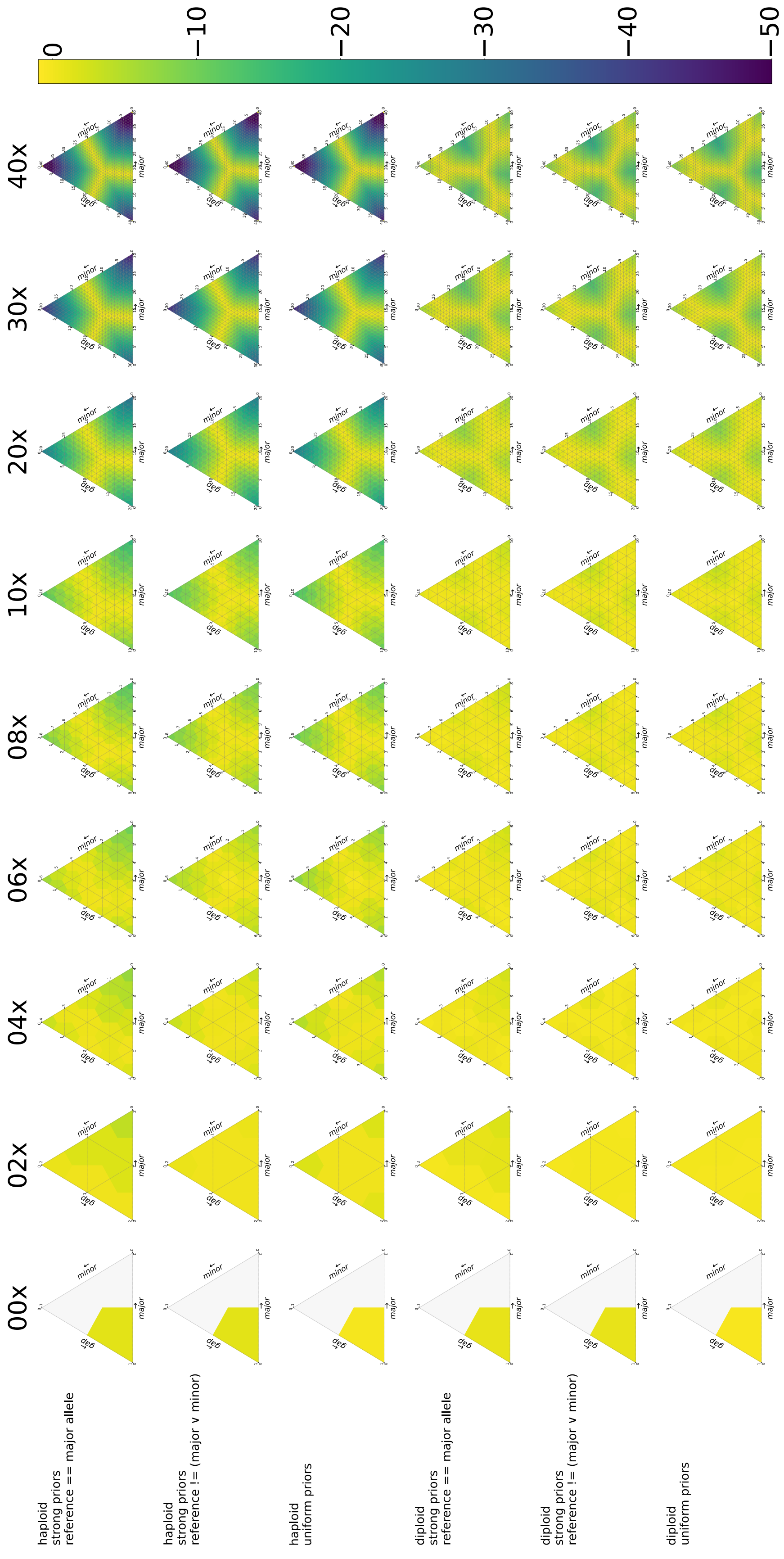

**Supplementary Figure 1: Uncertainty of various coverage patterns under different models.** We can express the observed coverage at a site as a combination of counts for a major (assumed to match reference, bottom edge), or minor (non-reference, right edge) allele, and counts for gaps (deletions, left edge). We plot log10 values of the uncertainty of the genotype posterior distribution given various coverage patterns. Each row of plots represents a different combination of model and priors, and each column is a combination of coverage counts summing to the value above the column. The models shown here comprise genotypes for either haploid or diploid genomes and are combined with either strong priors for the reference base or uniform priors across all possible bases (or gap). The second and fifth row additionally represent the cases where the observed bases do not include the reference base, i.e. a SNP. For the haploid models the uncertainty becomes lower when the coverage increases and the mixture moves closer to the vertices of the triangle. In contrast, the diploid model has additional zones of low uncertainty centered around the edge midpoints, which represent heterozygous genotypes. Notably, the choice of priors only affects uncertainty at low coverage levels. For example, uncertainty decreases faster when strong priors for the reference are used and most observed bases correspond to that nucleotide.

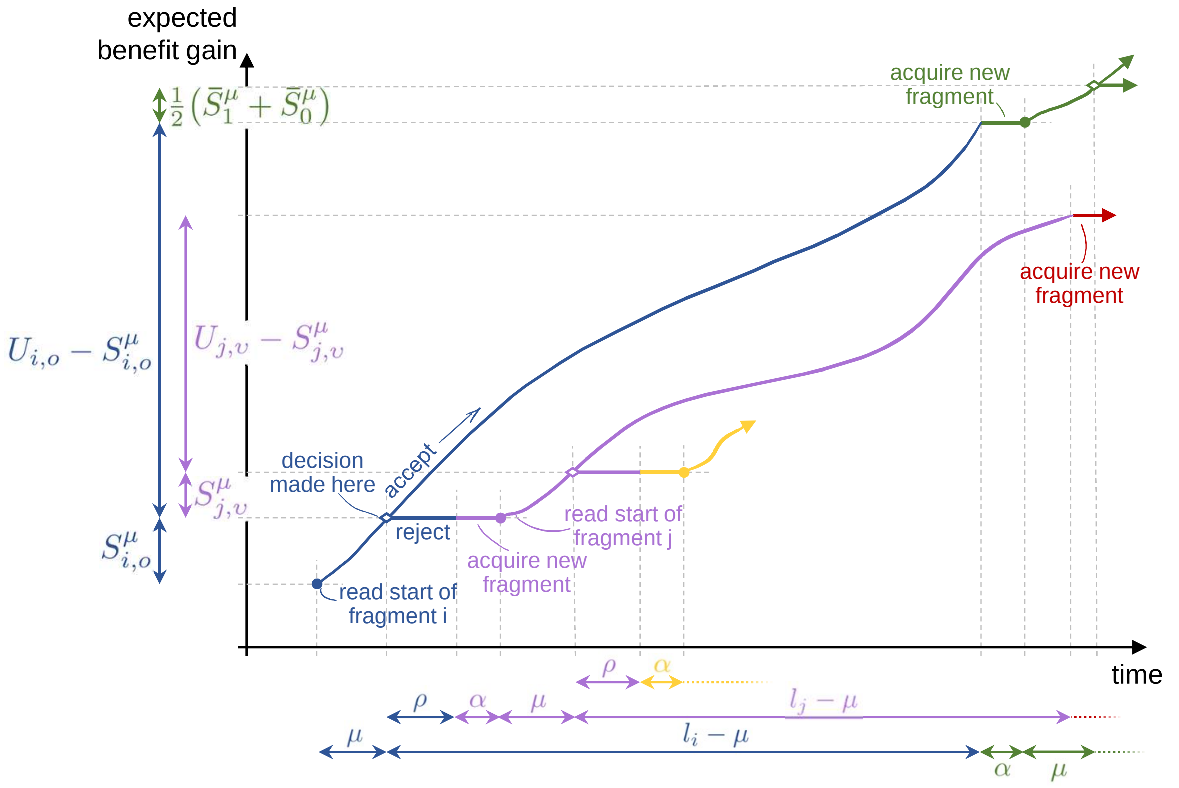

**Supplementary Figure 2: Schematic plot of our model of accumulated benefit against sequencing time.** Benefit gain is shown on the $y$-axis. For simplicity, we use an unrealistic scale for time on the $x$-axis. Colors show contributions of different DNA fragments. The process begins by a pore acquiring a fragment (blue). After time $\mu$, its origin and orientation $(i, o)$ are determined, benefit $S^{\mu}_{i,o}$ recorded, and the decision made whether to read the remainder of the fragment. If so, time $l_i - \mu$ passes (with $l_i$ the read length) generating further benefit $U_{i,o} - S^{\mu}_{i,o}$, after which a new DNA fragment is acquired (green, taking time $\alpha$) and mapped ($\mu$), and with expected benefit, prior to determination of position and orientation, of $(\bar{S}^{\mu}_1 + \bar{S}^{\mu}_0)/2$. Alternatively, the (blue) fragment can be rejected (taking time $\rho$; no further benefit) and a new fragment acquired (mauve; additional time $\alpha$). This in turn gets mapped ($\mu$; determining its location $j$, orientation $\upsilon$, and benefit $S^{\mu}_{j,\upsilon}$) and decided upon. Initial effects of other fragments are shown in gold and red. Filled circles mark points where new fragments are acquired; decision points are marked by open diamonds.

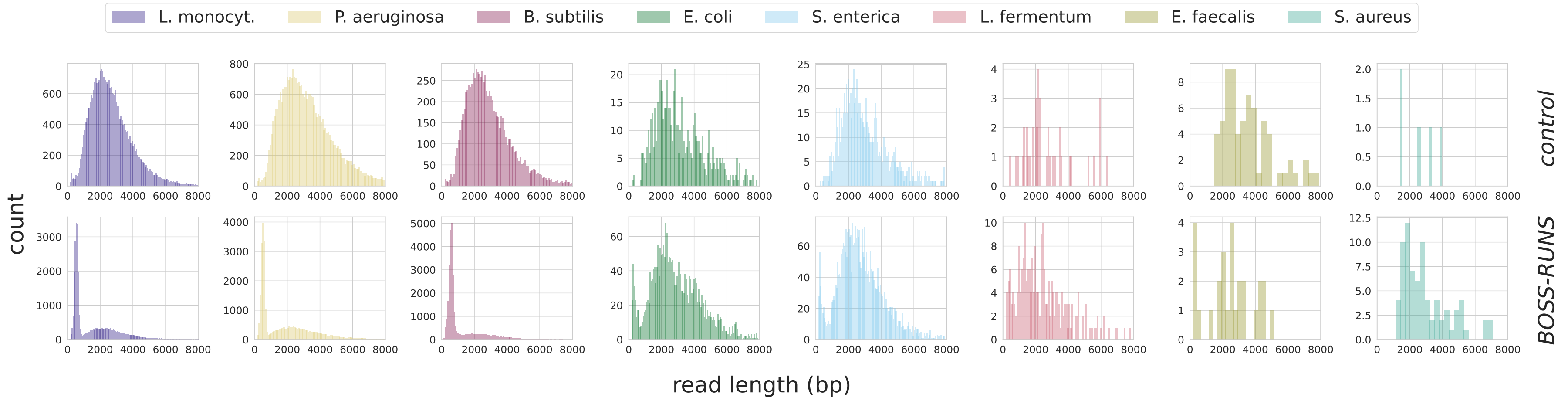

**Supplementary Figure 3: Distribution of read lengths separated by condition and source species.** In the main text, we show the overall distribution of read lengths for data from control channels and BOSS-RUNS. We further separated these distributions by the source species to check for potential differences in read lengths (at most 30k reads per species; top: control; bottom: BOSS-RUNS). Whereas we did not observe obvious differences between fragments from different bacterial species, we noticed that the peak at ∼450bp composed of rejected reads was visible even for some species where we did not expect to eject reads. The vast majority of these false negative decisions is caused by an inability to map the DNA fragment given its initial ($\mu$) bases. Increasing the amount of data used for the decision process could decrease the number of false rejections but would in turn decrease the advantage of rejection relative to reading the entire fragments. An alternative approach would be for reads of unknown origin to always be sequenced fully, which might be appropriate given prior knowledge about the input DNA.

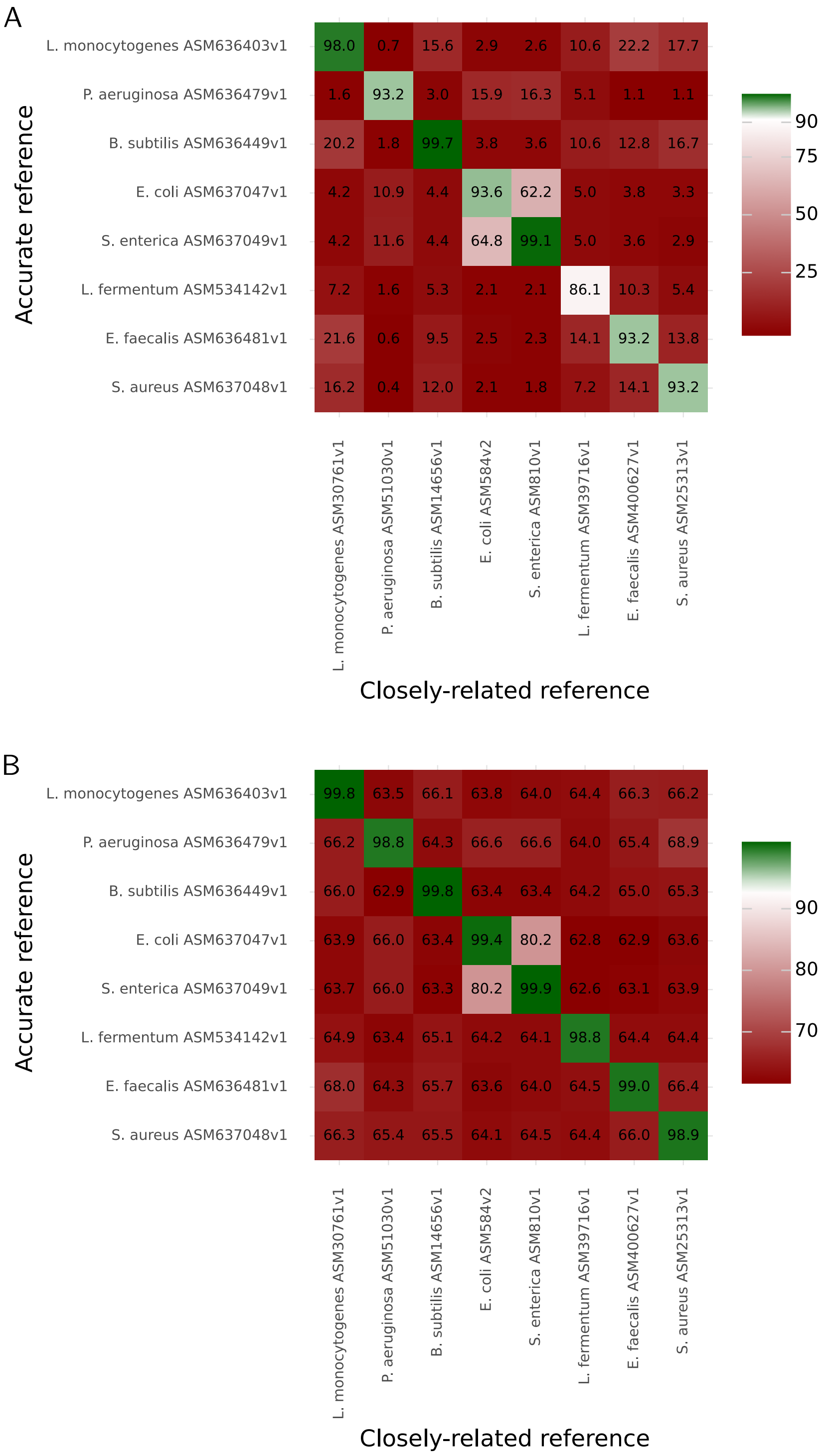

**Supplementary Figure 4: Divergence between bacterial strains in mock community and reference genomes used.** In order to mimic a realistic sequencing scenario, in which we do not have prior knowledge about the exact strains contained in a mixture, and to be able to analyse the ability to detect differences from the resulting data, we used references that are closely related but not identical to the true genomes. To quantify the difference we used both (A) the percentage of aligned nucleotide stretches (>90% sequence identity) between the true and reference assemblies and (B) ANI (average nucleotide identity) values determined using JSpecies in blast mode (Richter et al. 2016). The references used in our experiment are ordered along the $x$-axis by their abundance in the microbial community, while highly accurate assemblies (McIntyre et al. 2019) are shown along the $y$-axis. The color scale used transitions from red to green with a saddle point (white) at 0.9 in order to emphasize differences at high values.

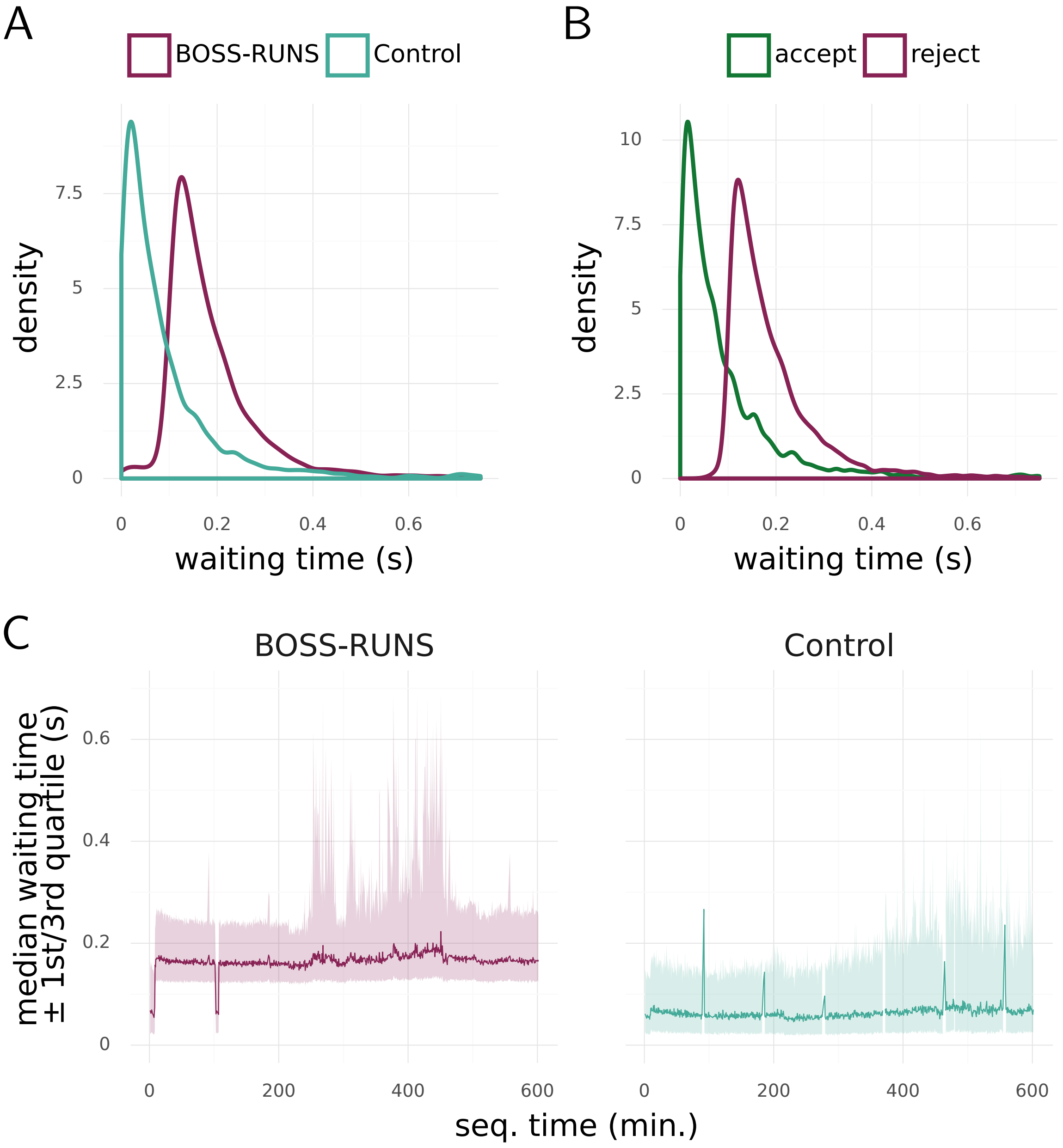

**Supplementary Figure 5: Analysing occupancy of pores during the sequencing experiment.** A) We quantified the elapsed time between data from two consecutive sequencing reads being transferred by the same pore on the flowcell. Separating all reads by condition shows that by rejecting reads the average time between reads in the same pore increases by about 100ms. This is caused by the time (modelled as $\rho$) taken to reject reads with BOSS-RUNS, with the BOSS-RUNS curve also showing the small proportion of accepted reads at far left. B) The waiting time distributions dependent on the executed decision, but irrespective of the condition, looks very similar to the previous plot. This further underlines that BOSS-RUNS successfully rejects most of the reads encountered. C) Throughout the sequencing experiment both the average waiting time and its variance increase slightly. This could be explained by multiple factors such as reduced pore health and depletion of fragments available for sequencing.

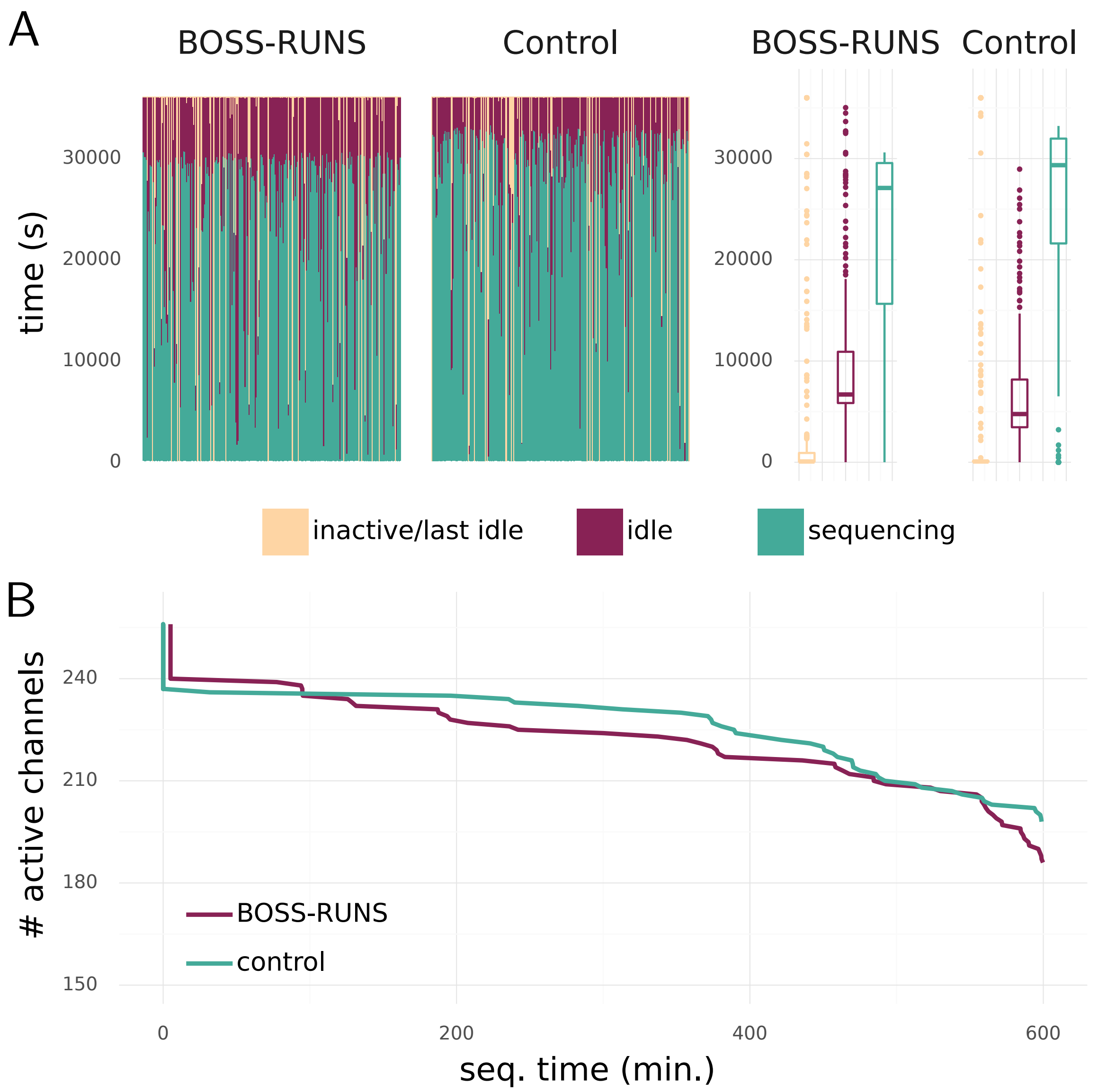

**Supplementary Figure 6: State of channels during the sequencing experiment** A) Left: Barplot showing the cumulative time of each channel spent sequencing, idling (i.e. rejecting a fragment or waiting for another one) and inactive (i.e. the final period not transmitting data before the end of the experiment). Right: Boxplots summarising these results across channels show that there is minimal difference between the time spent in each state. Inactive channels include those associated with blocked or damaged pores, which account for the long tails of outliers in the grey boxplots. B) The number of actively transmitting channels over time shows that using BOSS-RUNS does not significantly impact the rate of flowcell degradation. Active channels are all those that are not inactive (see A). A similar number of channels assigned to both methods were inactive before the experiment started, indicated by the initial drops in both lines shown at $t = 0$.

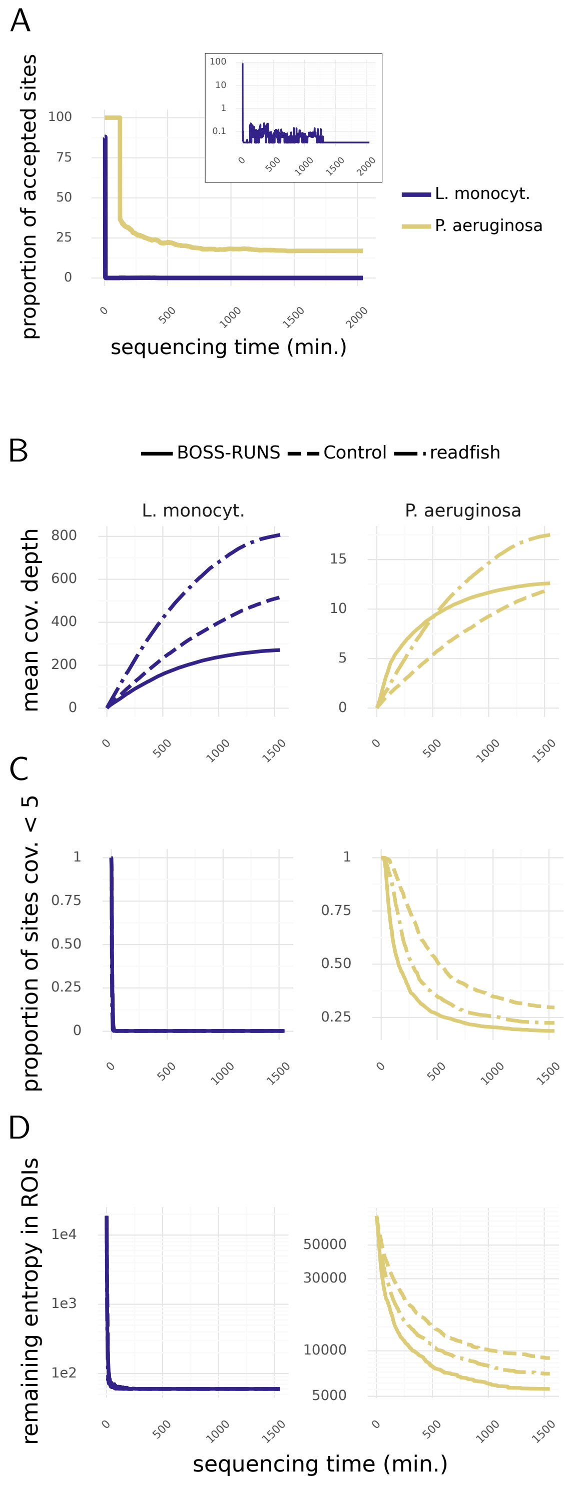

**Supplementary Figure 7: BOSS-RUNS mitigates coverage bias and redistributes coverage to sites of highest uncertainty when targeting ROIs.** A) While the proportion of accepted sites in *L. monocytogenes* is reduced very quickly, data from *P. aeruginosa* is sampled for longer and reaches a plateau after some time. The zoomed-in view reveals that individual sites of *L. monocytogenes* are still of interest during the experiment despite the order of magnitude difference in abundance. B) By sacrificing most of the coverage from the more abundant species, our method boosts coverage of the less-abundant bacteria. After some time, it too is mostly resolved and focusing on few, unresolved sites leads to lower average coverage compared to the control and readfish. C and D) In turn, the redistribution of coverage to sites where it is needed the most leads to both a lower proportion of sites at a coverage of $<5\times$ and a lower number of remaining unresolved sites (i.e. with highest posterior genotype probability $<0.99$.)

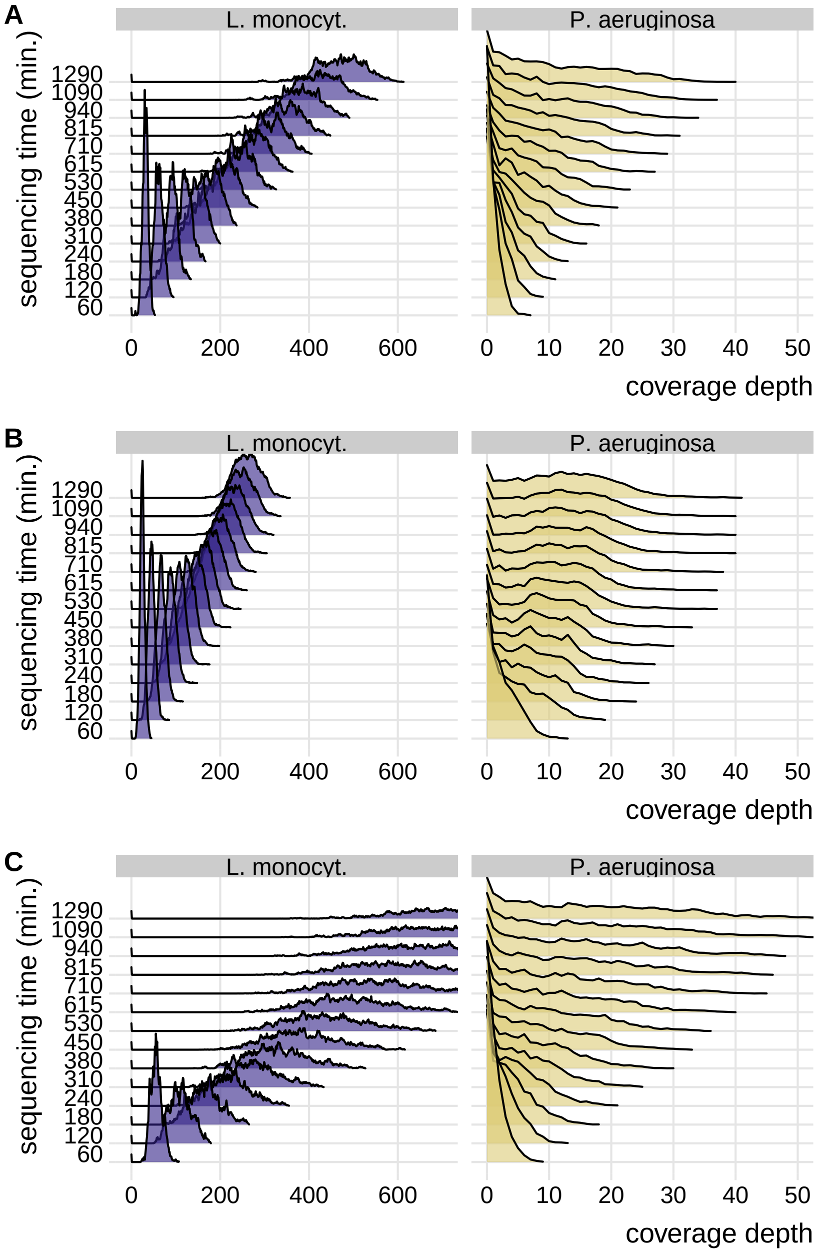

**Supplementary Figure 8: Coverage distribution of ROIs throughout the sequencing run.** Compared to the control sector of the flowcell (A), BOSS-RUNS (B) achieves lower variance of coverage depth in both species and ends up with fewer sites with low coverage ($<5\times$). Readfish (C) on the other hand continuously enriches for the *a priori*-specified ROIs and therefore has the highest levels of coverage in both bacteria. However, the number of remaining low coverage sites is larger.

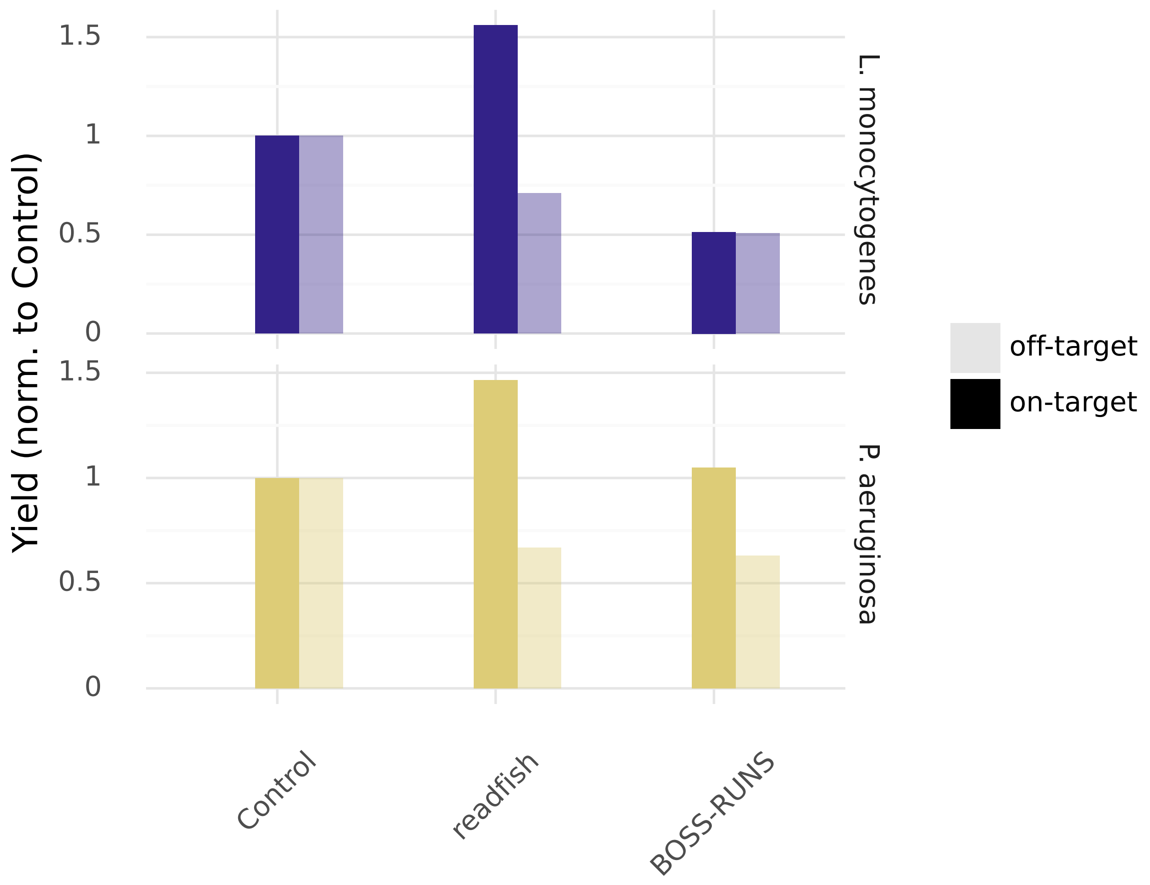

**Supplementary Figure 9: Enrichment of on-target and depletion of off-target regions compared to control.** By normalizing the acquired yield to the amount of data sequenced by the control condition, we can see that readfish successfully enriches for ROIs in both species and depletes sequence from the remaining genomes. Since BOSS-RUNS dynamically changes its decisions to focus on sites of highest uncertainty, it depletes both on- and off-target regions in *L. monocytogenes*, and only slightly increases yield of on-target regions in *P. aeruginosa*. However, BOSS-RUNS achieves its advantages by redistributing the sampled data within the genomes, which is not evident from solely looking at enrichment of ROIs.

### 4 Supplementary tables

**Supplementary Table 1:** Parameters and variables used in the BOSS-RUNS model

| | **Description** |
|---|---|
| $N$ | Reference genome size |
| $B$ | Set of observable characters (bases).<br>e.g. for haploid genomes without deletions $B = \{\mathrm{A}, \mathrm{C}, \mathrm{G}, \mathrm{T}\}$ |
| $G$ | Set of observable genotypes for the sequenced genome |
| $\theta$ | Prior probability for a substitution relative to the reference genome |
| $p_{\text{homo}}$ | Prior prop. of homozygous sites different from reference in a diploid genome |
| $b_R$ | Reference genome nucleotide at a position |
| $e$ | Probability that a nucleotide is mis-read as a different nucleotide |
| $L(l)$ | Probability that a fragment has length $l$ |
| $\eta$ | Number of values used to approximate $\widetilde{CL}(l)$ |
| $\rho$ | Time required to reject a fragment |
| $\alpha$ | Time required to acquire a new fragment |
| $\mu$ | Length used to map a fragment |
| $\mathcal{S}$ | Dynamic, adaptive decision strategy |
| $I_{i,o}^{\mathcal{S}}$ | Decision function of strategy $\mathcal{S}$ for a fragment starting at $i$ with orientation $o$ |
| $F_{i,o}$ | Probability that a fragment starts at $i$ and has orientation $o$ |
| $\pi_i(g)$ | Prior probability of genotype $g \in G$ at position $i$ |
| $f_i(g\|D)$ | Posterior probability of genotype $g \in G$ at position $i$ given data $D$ |
| $\phi(d\|g)$ | Probability of character $d$ with genotype $g$ |
| $P(d\|D)$ | Posterior probability of sequencing character $d$ given data $D$ |
| $S_i$ | Benefit score from an additional sequence covering position $i$ |
| $S_{i,o}^{l}$ | Cumulative score of read starting in position $i$, orientation $o$ and length $l$ |
| $U_{i,o}$ | Expected benefit of a read starting in position $i$ and orientation $o$ |
| $\widetilde{CL}(l)$ | Complementary cumulative distribution of read lengths |
| $\mathcal{D}_L$ | Domain of $L$ (values of $l$ where the distribution is strictly positive) |
| $\lambda$ | Mean fragment length |
| $\widehat{\mathcal{S}}$ | Strategy with optimal rate of benefit gain |
| $U_{i,o}^{\mathcal{S}}$ | Expected benefit of a fragment at $i$ with orientation $o$ under strategy $\mathcal{S}$ |
| $t_{i,o}^{\mathcal{S}}$ | Expected cost of a fragment at $i$ with orientation $o$ under strategy $\mathcal{S}$ |
| $\bar{U}^{\mathcal{S}}$ | Expected benefit of strategy $\mathcal{S}$ for next fragment |
| $\bar{t}^{\mathcal{S}}$ | Expected cost of strategy $\mathcal{S}$ for next fragment |
| $\bar{S}_o^{\mu}$ | Expected benefit of a read of length $\mu$ and orientation $o$ |

**Supplementary Table 2:** Percentage of sites with coverage $>5\times$ in regions flagged by Repeat-Masker

| Species | in masked region | not in masked region |
|---|---|---|
| *E. coli* | 2.191 | 97.809 |
| *S. enterica* | 0.180 | 99.820 |
| *L. fermentum* | 0.000 | 100.000 |
| *S. aureus* | 0.000 | 100.000 |
